# Musashi1 and its short C-terminal variants regulate pluripotency states in embryonic stem cells

**DOI:** 10.1101/2022.05.02.490263

**Authors:** Youwei Chen, Hailin Zhang, Jiazhen Han, Qianyan Li, Ying Chen, Gufa Lin

## Abstract

Musashi1 (MSI1) is a marker for adult stem cells, but little is known for its expression and function in pluripotent stem cells (PSCs). Here we report that MSI1 is expressed in embryonic stem cells (ESCs) and is required for pluripotency maintenance. We found that there exit short c-terminal MSI1 variants (MSI1-C, hMSI1^272-362^ or mMSI1^138-362^) in naïve but not primed ESCs. When overexpressed, MSI1 and MSI1-C variants facilitate primed-to-naïve pluripotency transition by elevating the pluripotency of primed hESCs toward a formative-like state, enable better survival of hESCs in human-mouse interspecies cell completion, and enhance the ability of blastoid formation of hESCs after naïve induction. Mechanistically, we show that the MSI1-C variants, though lacking RNA recognition motifs, bind to RNAs, enhance stress resistance and upregulate DNA damage repair genes. Thus, this study demonstrates that ESCs utilize MSI1 and the newly identified short MSI1-C proteins as double swords to regulate pluripotency states.

## INTRODUCTION

Musashi1 (Msi1) is an RNA-binding protein that has versatile functions in adult stem cells and cancer stem cells(Fox *et al*, 2015). It regulates proliferation and differentiation of neural stem cells (NSCs) and intestine epithelial stem cells (Cambuli *et al*, 2015; Good *et al*, 1998). Our current understanding of the mechanism of Msi1 has mostly derived from studies related to its RRMs (RNA recognition motifs). Msi1 contains two RRMs at the N terminal and is highly conserved among species(Sakakibara *et al*, 1996; Shibata *et al*, 2012; Yoda *et al*, 2000). The loss-of-function mouse models were generated by deleting the N-terminal sequences involving the RRM1(Sakakibara *et al*, 2002). Structural analysis of the binding activities of RRMs showed that the RMM1 domain is responsible for most of its target binding (Ohyama *et al*, 2012). There has been little attention on the C-terminal sequences of MSI1. This is largely due to the intrinsically disordered regions (IDRs), of which the structure is difficult to determine.

Recent evidence suggests that the C-terminal domain of MSI1 may also interact with RNAs(Chen & Huang, 2020; Chiou *et al*, 2017; Wang *et al*, 2020a), but the biological functions, if any, of the potential binding, has not been adequately investigated.

On the other hand, while most studies of MSI1 have focused on its roles in adult stem cells, little is known about its expression and function in pluripotent stem cells (PSCs), which may exist in one of the three pluripotency states (Kinoshita *et al*, 2021; Smith, 2017). PSCs can exist in the formative state, in addition to the naïve and primed pluripotency states. The state PSCs reside in have great impact in term of the proliferation and differential potentials. Both PSCs in the naïve and the primed states can differentiate into cells of all three germ layers in culture, or form teratomas when implanted(Lee *et al*, 2017). But only naïve PSCs can contribute to germline chimeras (Eggan *et al*, 2002; Zhao *et al*, 2009). Primed PSCs cannot form chimeric embryos efficiently(Huang *et al*, 2018). Recent works showed that PSCs in the formative state have the higher potential to contribute to chimeric embryonic development than the primed state PSCs (De Los Angeles & Wu, 2021; Yu *et al*, 2021b). Thus, a full understanding of pluripotent states regulation is critical for obtaining formative and naïve pluripotency states to pave the way for translational applications of human PSCs, including human embryonic stem cells (hESCs) (Yu *et al*., 2021b). Yet hESCs in cultures are considered as in a primed pluripotency state, which has disadvantages in expansion and do not contribute to chimeras. Therefore, there has been much effort to obtain naïve and formative human PSCs(Chan *et al*, 2013; Gafni *et al*, 2013; Theunissen *et al*, 2014). However, when converted, human naïve PSCs still suffer from low genomic stability(Kilens *et al*, 2018; Yu *et al*, 2021a).

In our analysis of the MSI1 regulated RNAs during PSC derived NSCs, we observed that MSI1 is expressed in R1 mESCs and H9 hESCs. This raised the possibility that MSI1 also functions in PSCs. We found that loss-of-function of *Msi1* in R1 mESCs caused pluripotency dissolution, indicating that MSI1 regulate pluripotency in mESCs. Interestingly, MSI1 is distinctly localized in the mouse versus human ESCs, and there exit short C-terminal MSI1 variants (MSI1-C) in the naïve R1 and naïve induced H9 ESCs, but not in the primed state H9 cells. When over-expressed, MSI1 and the RRM truncated MSI1-C variants enhance the pluripotency of primed hESCs toward a formative-like state, increasing hESC survival in interspecies cell competition. More strikingly, MSI-C variants could also prolong blastoid formation capacity of naïve induced hESCs. Thus, our result show that MSI1 regulates pluripotency states with the full-length and the short MSI1-C variants. Mechanistically, we found that both MSI1 and the C-terminal MSI1-C proteins binds to RNAs, enhance stress resistance and upregulate DNA repair genes in hESCs during naïve state acquisition, leading to better survival of hESCs.

## RESULTS

### MSI1 is expressed in ESCs with distinct subcellular localization between primed and naïve states

In our analysis of MSI1 during NSC differentiation from H9 hESCs, we found that in addition to its upregulation in NSCs, MSI1 is expressed in hESCs (Figure S1A, Figure 1A). While its function in embryonic neurogenesis is well established (Sakakibara *et al*., 1996; Sakakibara *et al*., 2002), the role of Msi1 in ESCs has not been explored. RNA-immunoprecipitation-PCR (RIP-PCR, with anti-MSI1, abcam ab52865) assay showed that in addition to binding to *Sox1, CDKN1A* and *CDK6* in NSCs, MSI1 binds to pluripotency markers *POU5F1, NANOG* and *SOX2* in H9 hESCs, suggesting a possible role of MSI1 in ESCs (Figure S1B). We also examined MSI1 expression in R1 mESCs and found that it is expressed, but unlike the cytoplasmic expression of MSI1 in hESCs, MSI1 signals are localized in the nucleus (Figure 1A). Western blot analysis with fractionated nuclear and cytoplasmic extracts confirmed the distinct subcellular localization of MSI1, but not its homolog MSI2, in H9 versus R1 ESCs (Figure 1B).

**Fig. 1.**
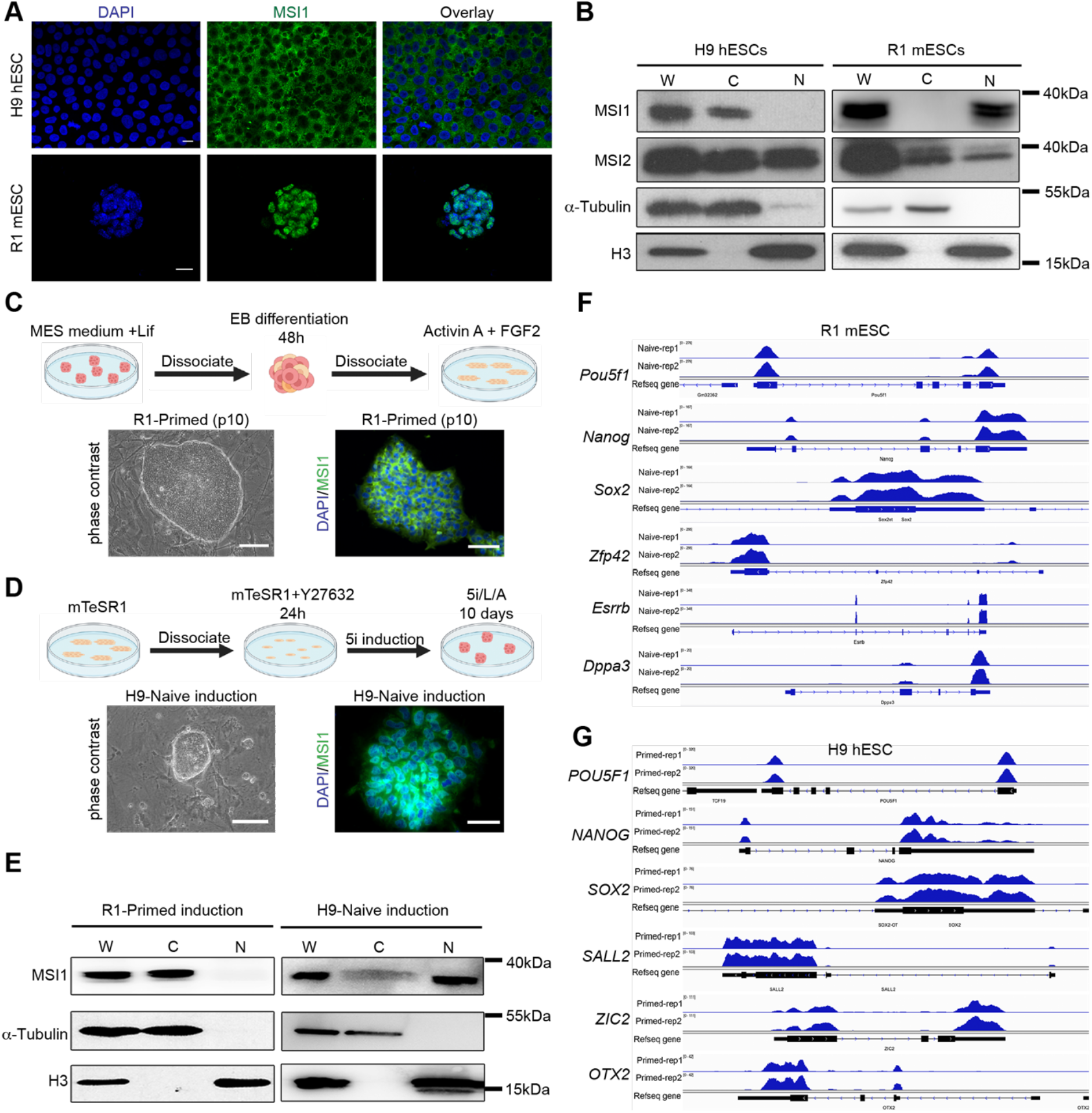
MSI1 is expressed in primed and naïve state ESCs with distinct subcellular localization. (**A**) Immunofluorescence (IF) detection of MSI1 (green) in H9 hESCs and R1 mESCs, with anti-MSI1 antibody (ab52865), showing cytoplasmic versus nuclear localization of MSI1. Nuclei were stained with DAPI. Scale bar, 20 μm. (**B**) Western blot detection of MSI1 and MSI2 (ab76148) with fractionated protein extracts in H9 hESCs and R1 mESCs. α-Tubulin is used as loading control for cytoplasmic proteins, and Histone H3 for nuclear proteins. (**C**) Induction of R1 mESCs to primed state, and immunofluorescence staining of MSI1 showing MSI1 expression in the cytoplasm of passage 10 R1-primed cells. Scale bar, 100 μm (**D**) Induction of H9 hESCs to naïve state and detection of MSI1 by immunofluorescence on P5 naïve induced H9 cells (on Matrigel for staining). Scale bar, 100 μm (**E**) Western blot detection of MSI1 with fractionated extracts in R1-primed mESCs and naïve induced H9 hESCs. α-Tubulin is used as loading control for cytoplasmic proteins, and Histone H3 for nuclear proteins. (**F**) Integrative Genomics Viewer (IGV) plots of pluripotency genes *Pou5f1, Nanog, Sox2*, and naïve state genes *Zfp42, Esrrb, Dppax3* in R1 mESCs, with position (x-axis) and count (y-axis) overlapped with MSI1 RIP-Seq reads. (**G**) IGV plots of *POU5F1, NANOG, SOX2* and primed state genes *SALL2, ZIC2, OTX2* in H9 hESCs, with position (x-axis) and count (y-axis) overlapped with MSI1 RIP-Seq reads.

To test whether the distinct localization of MSI1 is related to the pluripotency states of mESCs and hESCs (in the naïve and the primed state, respectively)(Huang *et al*., 2018), we examined MSI1 expression in the mouse embryos, at E3.5 (before embryo implantation) and E5.5 (after embryo implantation) of development. MSI1 signals were mainly localized in the nucleus of E3.5 embryo. But in the E5.5 embryo, MSI1 signals were mainly found in the epiblast, though detectable in the extra embryonic ectoderm (Figure S1C). To analyze whether the observed dynamic expression of MSI1 is related to the transition of naïve and primed pluripotency, we induced R1 mESCs to a primed state by FGF2/Activin A (Zhang *et al*, 2010). The induced R1 cells (R1-primed, passage 10, p10) were positive for primed state markers FGF5 and T, with maintained expression of pluripotency markers NANOG, OCT4 (POU5F1) (Figure S1D,E).

Interestingly, the induced R1-primed cells showed cytoplasmic expression of MSI1(Figure 1C, E). Then we used the 5i system(Theunissen *et al*., 2014) to convert H9 hESCs from a primed state to a naïve state (H9-5i). The H9-5i colonies were dense and spherical, with shiny edges (Figure 1D). The H9-5i cells had high levels of *OCT4* (*POU5F1*), *NANOG*, and the naïve state markers *ZFP42* (*REX1*), *DNMT3L* and *TBX3* (Figure S1F). Both immunofluorescence and Western blot analysis showed that MSI1 appeared in the nuclei of H9-5i cells, suggesting that naïve induction induced nuclear expression of MSI1(Figure 1D,E).

RIP-seq (with anti-MSI1, ab52865, isotype IgG as control) showed that MSI1 binds to transcripts of pluripotency markers *POU5F1, NANOG*, and *SOX2* in both R1 and H9 ESCs. In addition, MSI1 binds to RNAs of the naïve marker genes *Zfp42* (*Rex1*), *Esrrb*, and *Dppa3* in R1 mESCs, while in the H9 ESCs it binds to the RNAs of the primed markers *OTX2, ZIC2*, and *SALL2* (Figure 1F,G). Thus, these results showed that the subcellular distribution of MSI1 is related to the pluripotency states of ESCs.

### MSI1 is required for pluripotency maintenance in R1 mESCs

To investigate the function of MSI1 in ESCs, we used CRISPR-Cas9 system to knock out *MSI1* in the R1 mESCs and H9 hESCs. We designed two gRNAs targeting exon 1 and exon 9 of the mouse *Msi1* (*mMsi1*) gene (all known and predicted *Msi1* transcripts are covered by the gRNAs, www.ncbi.nlm.nih.gov) (Figure 2A), and obtained two *mMsi1* deletion cell lines, R1-C3 and R1-C5. Sequencing results verified deletions of the *mMsi1* gene (exon1-9) in both lines, and we used R1-C5 for further analysis (Figure 2B). Western blot analysis confirmed the deletion of MSI1 protein in the R1-C5 cells (Figure 2C). While the R1-C5 clone could be expanded as single cell, round shaped cells gradually emigrated from the spherical colonies (Figure 2D, asterisks). Although expandable by single cell passaging (which is routinely used for mouse ESCs cultured on feeder cells) for a couple of generations, R1-C5 cells eventually failed to form colonies (Figure 2E). We detected the levels of 5mC and 5hmC in the genomic DNA isolated from the R1-C5 colonies (differentiated cells depleted, see method) and found that the still expandable R1-C5 colonies have higher levels of DNA methylation (Figure 2F,G). Immunofluorescence analysis showed that the still expandable R1-C5 cells have lost expression of the naïve pluripotency marker Rex1 (ZFP42) and started to express the primed pluripotency marker T (Figure S2A). RT-qPCR of the R1-C5 colonies (differentiated cells depleted, see method) showed that the R1-C5 cells have lower expression of *Pou5f1, Nanog* and *Sox2*, and higher expression of ectoderm, mesoderm and endoderm associated marker genes, with *Gata4, Foxa2* and *Gata6* being the most up-regulated (Figure S2B). RNA-seq analysis of R1-C5 and R1-WT cells also confirmed the down regulation of gene involved in the naïve pluripotency regulation and upregulation of genes involved in differentiation toward endoderm (Figure 2H, Figure S2C-E). By overexpression of Msi1 in the R1-C5 cells, we observed that this pluripotency withdrawal and differentiation can be rescued (Figure 2I). Thus, these observations showed that deletion of MSI1 in the naïve R1 mESCs causes pluripotency dissolution.

**Fig. 2.**
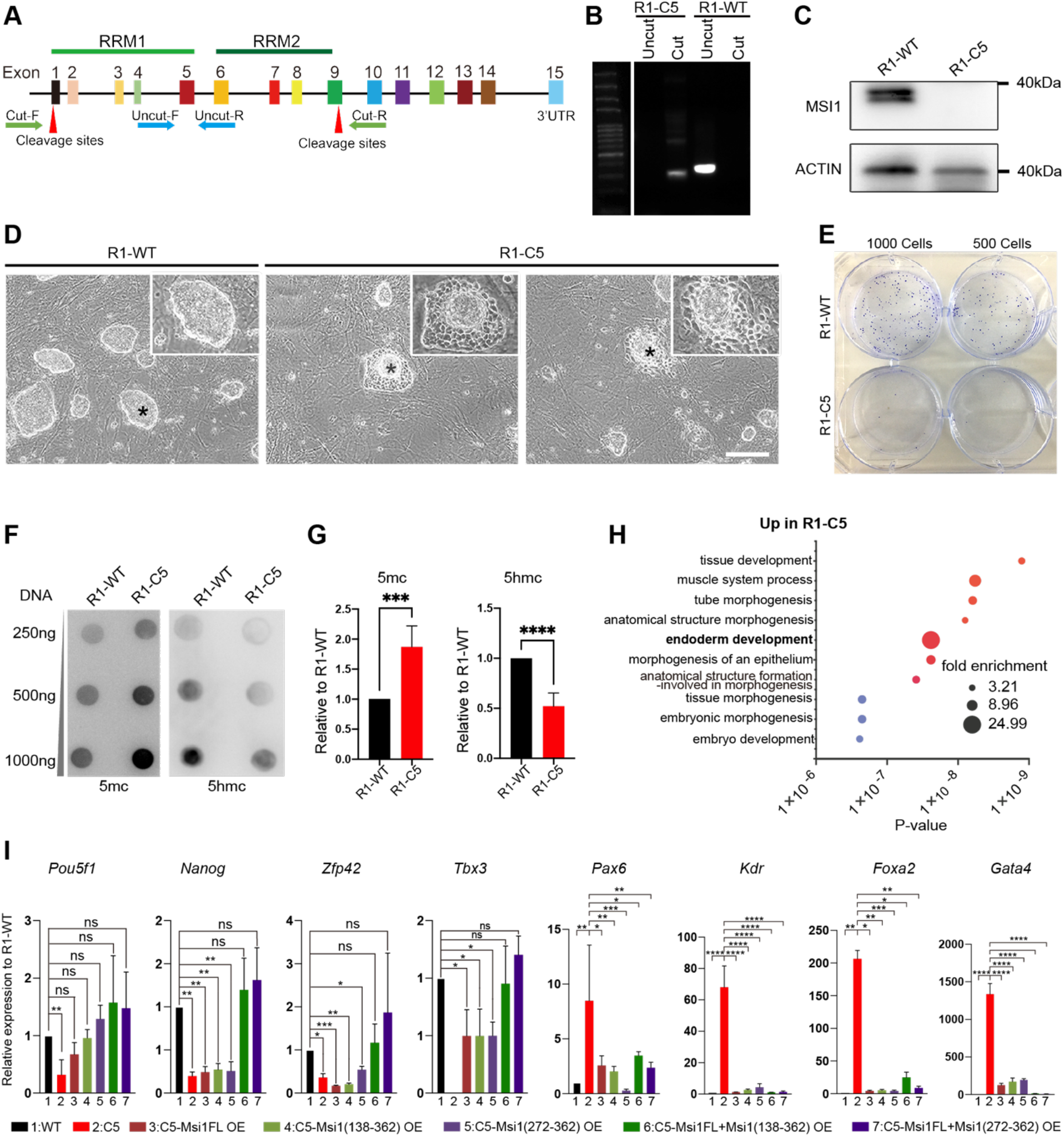
Requirement of MSI1 for pluripotency maintenance in mESCs. (**A**) Schematic diagram of CRISPR-Cas9 deletion of mouse *Msi1* gene in R1 mESC. The green arrows show positions of primers for deletion detection, the blue arrows for non-deletion detection, the red arrows represent the Cas9 protein cleavage sites. (**B**) Genomic PCR detection of R1-C5, a clone with expected deletion of mouse *Msi1*. (**C**) Western blot detection of MSI1 (with ab52865) in wild type R1 (R1-WT) and R1-C5 mESCs, showing absence of MSI1 expression in R1-C5, a cell line for mMsi1 deletion. (**D**) Phase contrast micrographs of R1-WT and R1-C5 cells, showing emergence of differentiated cells around the R1-C5 colonies. Insets show higher magnifications of the colonies marked with *. (**E**) Crystalline violet staining of R1-WT and R1-C5 colonies for colony forming assay, with seeding of 1000 or 500 cells. (**F**,**G**) Analysis of 5mc and 5hmc levels in genomic DNA of R1-C5 and R1-WT cells, by dot-blotting assay (**F**), and intensity quantitation (**G**). (**H**) GO enrichment analysis for DEGs in R1-C5 cells against R1-WT cells, showing the top 10 up-regulated biological processes. (**I**) RT-qPCR detection of *Pou51f, Nanog, Zfp42, Tbx3, Pax6, Kdr, Foxa2, Gata4* in R1-WT and R1-C5 cells, with overexpression of *Msi1* and *Msi1-C* variants. Asterisks indicate P-values. (*, p<0.05, **, p<0.01, ***, p<0.001). ns, no significant difference. t-test, from 3 independent replicates. Scale bars (**D**), 100 μm.

### Short MSI1 C-terminal proteins are expressed in H9 hESCs after RMM truncation

We deleted human *MSI1* gene (h*MSI1*) in H9 hESCs by CRISPR/Cas9. When the exons 1-14 of the *hMSI1* were targeted, we did not obtain expandable clones (Table S1), demonstrating that MSI1 is also necessary for pluripotency maintenance in hESCs. Interestingly, when we targeted the exons 1-7 of *hMSI1* to delete the RRMs (Figure 3A), we obtained an expandable H9 hESC lines, H9-C8 (Figure S3A). This strategy was similar to the approach previously used to knockout MSI1 in animal models, in which the N-terminal RRM1 of Msi1 was deleted (Sakakibara *et al*., 2002). Western blot and immunofluorescence analysis (with anti-MSI1, ab52865) showed that H9-C8 cells did not have positive signals (Figure 3B, Figure S3B). However, unlike R1-C5 cells, H9-C8 cells maintained the typical colony morphology during passaging, and there was no sign of cell differentiation (Figure 3C). Moreover, RT-qPCR results showed that H9-C8 cells have similar expression levels of *POU5F1, NANOG, SOX2, TBX3, DNMT3L* and *ZFP42*, except for a slightly lower level of *KLF4* (Figure S3C).

**Fig. 3.**
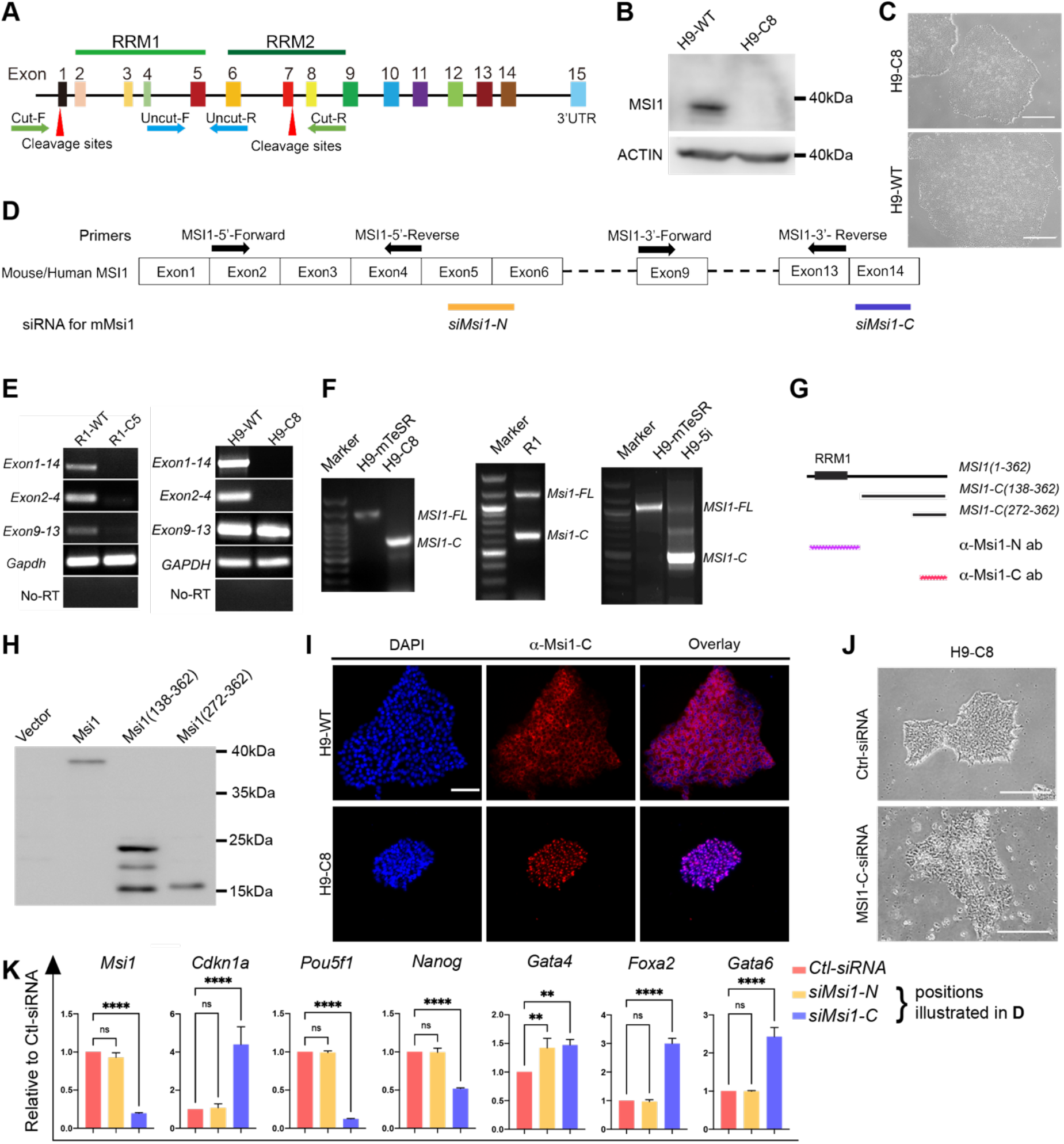
Identification of MSI1-C variants in ESCs. (**A**) Schematic diagram of CRISPR-Cas9 deletion of human MSI1 gene in H9 hESC. The green arrows show positions of primers for deletion detection, the blue arrows for non-deletion detection, the red arrows represent the Cas9 protein cleavage sites. (**B**) Western blot detection of MSI1 (with ab52865) showing absence of MSI1 expression in H9-C8 hESCs. ACTIN as loading control. (**C**) Morphology of H9-C8 and H9-WT colonies. Scale bar, 100 μm. (**D**) Schematic diagram of primers used for detecting the 5’ N-terminal (Exon 2-4) and 3’ C-terminal (Exon 9-13) sequences of mouse and human MSI1 transcripts. Positions of siRNA against *mMsi1* related to (**K**) shown below the gene. (**E**) RT-PCR detection of Msi1 transcripts in R1-WT, R1-C5, H9 and H9-C8 ESCs. Note that transcripts from exon 9-13 was detected in H8-C9 hESCs. Exon 1-14 was detected by primers for the whole coding sequences. No-RT, no-reverse transcriptase control. (**F**) 5’RACE detection of MSI1 transcript variants in R1 and H9 ESCs. Short variants were sequenced to deduce the proteins encoded (mMSI1138-362 in R1; hMSI272-362 in H9-5i). (**G**) Illustration of the overexpression vectors used in H9 cells, and the positions of antigen domain for antibody production (from abcam, Gentex). (**H**) Western blot analysis of MSI1 and MSI1-C variants overexpressed in H9 hESCs, showing the translation of MSI1 and MSI1-C variants, detected by α-Flag antibody. Flag tags were added to both the N-and C-terminus of the overexpression constructs, to detect potential variants. Note MSI1138-362 can produce further shortened proteins. (**I**) Immunofluorescence with a-MSI1-C in H9-WT and H9-C8 hESCs. Note nuclear localization of a-MSI-C signals in H9-C8 cells. Scale bar, 100 μm. (**J**) Phase contrast micrographs of hESC, after 48 h of siRNA knockdown of the MSI1-C transcripts in in H9-C8 cells. MSI1-C-siRNA caused cell death and differentiation, while Control-siRNA had no effect. (**K**) RT-qPCR detection of Msi1-C, Cdkn1a, Pou5f1, Nanog, Gata4, Foxa2, Gata6 in R1 mESCs after transfection with siRNAs against the N- and C-terminal of mouse *Msi1* transcripts, with *siMsi1-N* and *siMsi1-C* (positions of the siRNA shown in **D**). Scale bars (**I-J**), 100 mm. * indicates p values (**, p<0.01; ***, p<0.001; ****, p<0.0001). ns, not significant. n = 3 in (**K**).

Previous studies showed the existence and expression of *Msi1* variants during newt retinal regeneration (Susaki *et al*, 2009). And analyzing the sequence of *hMSI1* and *mMsi1* genes implicated the presence of multiple open reading frames (ORFs), all of which end with a stop codon in the exon 14. We hypothesized that MSI1-C variants (MSI1-C), which is undetected by the ab52865 antibody (antigen sequence undefined) may be present in ESCs, when the full-length MSI1 is removed. Indeed, our series of analysis showed this is the case.

First, we designed PCR primers for the full-length, N-terminal (exon 2-4) and C-terminal (exon 9-13) of *MSI1* mRNAs (Figure 3D). RT-PCR results showed that while no *mMsi1* transcript exists in R1-C5 cells, primers against the C-terminal of *hMSI1* could amplify *hMSI1* transcripts in the H9-C8 cells (Figure 3E), suggesting that additional *hMSI1* transcripts are produced in H9-C8 cells, after deletion of the exons 1-7.

Next, we performed 5’RACE in the H9-WT and H9-C8 cells and found that H9-C8 cells lack full-length *hMsi1* transcript but do have short *hMsi1* transcript variant (Figure 3F). Sequencing of the short *hMsi1* transcript showed that it could encode a protein corresponding to hMSI1^272-362^ if translated (Figure S3D). This short transcript has never been reported.

Then, to verify that the short *hMSI1* transcripts can be translated, we constructed Flag tagged *MSI1-C* expression vectors (*Msi1*, full-length; *Msi1*^*272-362*^ and *Msi1*^*138-362*^, C-terminal MSI1) using *mMsi1* gene sequence (there is only one S/T difference at position 252 between mouse and human MSI1 proteins). We performed Western blot analysis with α-Flag, α-MSI1-N (ab21628, against aa1-100 of MSI1) and α-MSI1-C (GeneTex, GT2377, against C-terminal sequences of MSI1) antibodies. Judging from the antigen domains for producing the α-MSI1-N (ab21628) and α-MSI1-C (GT2377) antibodies (Figure 3G), we expected that α-MSI1-N would report protein band only in full-length *Msi1* transfected cells, while α-MSI1-C could report protein products in full-length *Msi1, Msi1*^*272-362*^ and *Msi1*^*138-362*^ transfected cells. Immunofluorescence and Western blot analysis confirmed the expected results, and thus the specificity of the N- and C-MSI1 antibodies (Figure S4A,B). As control, α-Flag reported protein products in all samples, showing similar Western blot bands to α-MSI1-C (Figure S4A,B). Transfection of H9-WT cells confirmed that *Msi1*^*272-362*^ could also be translated in H9-WT cells (Figure 3H).

Immunofluorescence analysis of the H9-C8 cells showed that H9-C8 cells, did not express full-length MSI1 proteins (Figure 3B), but did have nuclear signals detected by the α-MSI1-C antibody (Figure 3I), indicting the presence of MSI1-C proteins.

Last, to test whether the short MSI-C proteins are responsible for the maintained pluripotency in H9-C8 cells. We used siRNA against the C-terminal sequences of h*MSI1* in H9-C8 cells. Now we were left with a lot of dead cells and very few colonies (Figure 3J, Figure S3F).

### MSI1 C-terminal proteins are also present in R1 mESCs and naïve induced H9 hESCs

The nuclear localization of the MSI1-C proteins observed in H9-C8 cells raised the possibility that nuclear expression of MSI1 in R1 mESCs and naïve induced H9 cells (Figure 1) are also due to the existence of C-terminal MSI1-C proteins. Indeed, as in the H9-C8 cell, the 5’RACE results confirmed that upon 5i induction, new transcripts appeared in the H9-5i hESCs (Figure 3F). And in the naïve state R1 mESCs, there existed a short transcript in addition to the full-length m*Msi1* (Figure 3F). Sequencing of the 5’RACE products showed that the main short C-terminal transcript in H9-5i also encodes hMSI1^272-362^ but the short C-terminal transcript in R1 mESCs has an ORF encoding mMSI1^138-362^ (Figure S3E). Interestingly, in *MSI1*^*138-362*^ transfected cells, additional smaller western bands were present, indicating that further shortened MSI1-C variants (including, probably, MSI1^272-362^, the smallest band) were produced (Figure 3H, Figure S4B). Thus, these results showed that there exist short MSI1-C proteins, mMSI1^138-362^ in the naïve mESCs and hMSI1^272-362^ in naïve induced hESCs, in addition to the full-length MSI1.

### Msi1-C proteins increase efficiency of primed-to-naïve state transition

The existence of additional short MSI1-C proteins in the naïve R1 mESCs and the 5i induced H9 hESCs suggested that the MSI1-C proteins have a specific role in pluripotency state regulation. Consistent to the observed pluripotency withdrawal in R1- C5 cells, we found that in H9-WT cells, siRNAs against the N-or C-terminal of *mMsi1* (targeting positions to mMsi1 illustrated in Figure 3D) also decreased expression of pluripotency markers and increased expression of endoderm markers (Figure 3K). But the siRNA against *mMsi1* C-terminus was more effective. Moreover, overexpression of the *Msi1-C* variants, on the other hand, could reverse expression levels of the endoderm marker genes *Kdr, Gata4* and *Foxa2*, and the ectoderm marker gene *Pax6* in R1-C5 cells. Combined expression of full-length Msi1 and Msi1-C variants could also sufficiently rescue expression of pluripotency markers (Figure 2I).

Considering that MSI1-C proteins are expressed in naïve but not primed ESCs (Figure 3), we wondered whether MSI1-C variants also have a role in establishing naïve pluripotency, such as during the process of primed-to-naïve transition. First, we utilized H9-C8 cells for naïve induction, as H9-C8 cells have endogenously expressed MSI1-C proteins. Like H9, H9-C8 cells could be efficiently converted to naïve state and the converted H9-C8-5i cells have higher expression levels of *POU5F1, NANOG, KLF4, DNMT3L, TBX3*, though a similar level of *ZFP42*, compared to H9-WT-5i cells (Figure 4A). In addition, colony forming assay showed that H9-C8-5i cells can form more colonies than H9-WT-5i cells (Figure 4B). Then we overexpressed MSI1 and MSI-1^272-362^ in H9-WT cells and subjected them to 5i induction. RT-qPCR results showed that significant higher level of *NANOG, POU5F1, ZFP42* and *DNMT3L* was induced (Figure 4C). Thus, these observations indicated that MSI1-C proteins facilitate the establishment of naïve pluripotency.

**Fig. 4.**
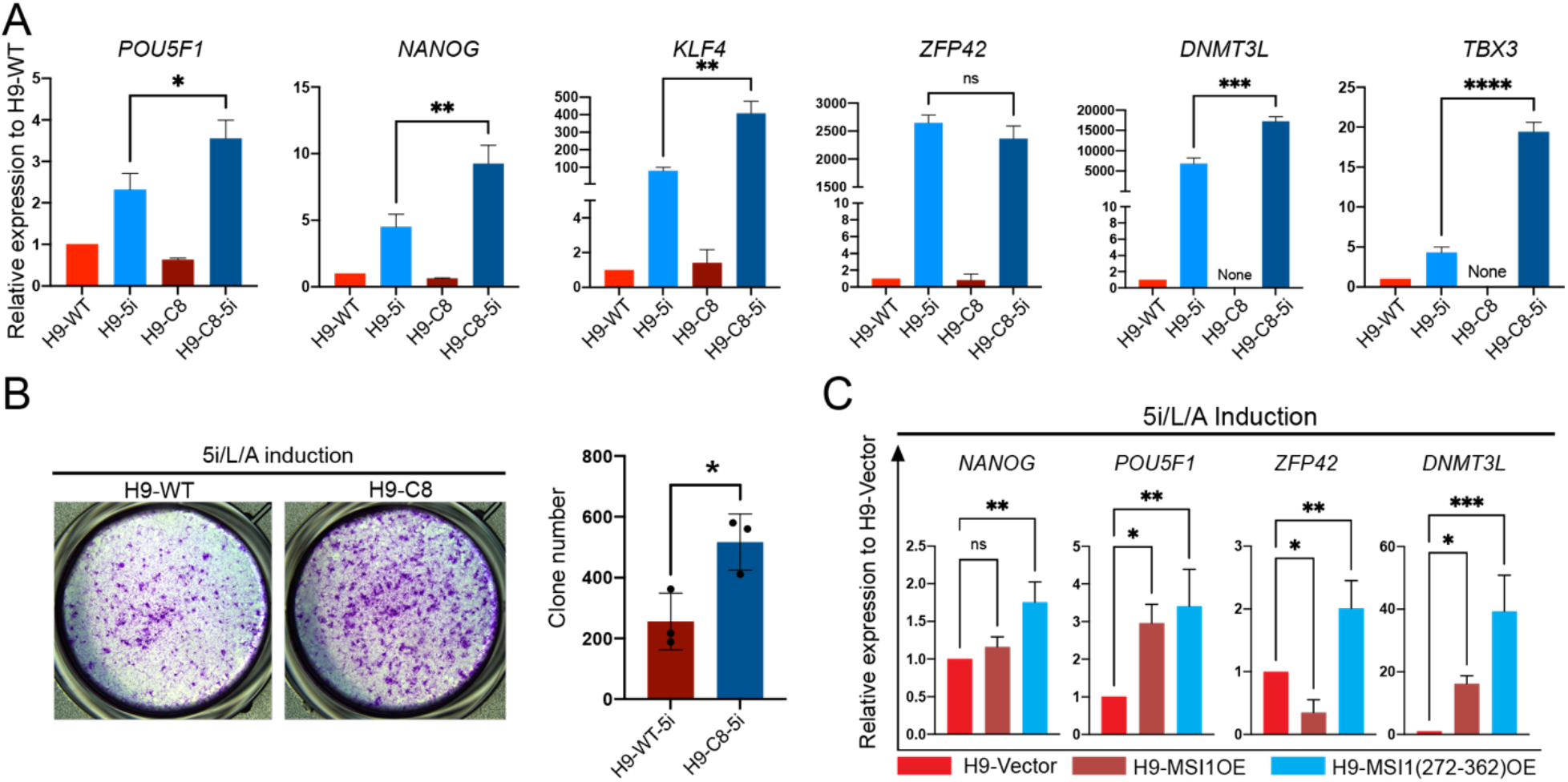
MSI1-C variants increase efficiency of primed-to-naïve state transition. (**A**) RT-qPCR detection of pluripotency genes *POU5F1, NANOG, KLF4* and naïve genes *ZFP42, DNMT3L, TBX3* in H9-WT and H9-C8 hESCs after 5i naïve induction. (**B**) Colony forming analysis of 5i induced H9-WT-5i and H9-C8-5i cells. (**C**) RT-qPCR detection of pluripotency genes *POU5F1, NANOG, ZFP42* and *DNMT3L* in H9-WT hESCs with overexpression of MSI1, MSI1-C variants, after 5i naïve induction. (**D**) indicate *P*-values (*, p<0.05; **, p<0.01; ***, p<0.001). ns, no significant difference. *t*-test, from 3 independent replicates.

### MSI1-C variants enhance stress response of hESCs with increased mitochondrial activity

To understand the underlying mechanism of MSI1-C proteins in facilitating pluripotency transition, we performed RNA-seq analysis to identify DEGs (differentially expressed genes) in H9-WT, H9-C8, H9 hESCs overexpressed with MSI1 or MSI1-C variants (H9-Msi1C-OE) (Supplementary File S2). There were 6654 DEGs commonly shared between MSI1 and MSI1-C OE vs H9 WT cells. GO analysis revealed that the most significantly up-regulated genes enriched biological processes were DNA double-strand break process, stress granule assembly and negative regulate stem cell differentiation (Figure 5A). As shown in the heatmap plot (Figure 5B), genes involved in the processes of stress granule assembly were highly upregulated in MSI1-C overexpressed H9 cells, as well as in H9-C8 cells. We identified 296 DEGs between H9-C8 (which express Msi1-C) vs. H9-WT cells, and found that the most upregulated processes in H9-C8 cells (which express MSI1-C proteins) were all related to stress responses and detoxification of metal ions, too (Figure 5C).This strongly suggested that MSI1-C proteins are involved in stress response, which is in consistence with the ability of stress granule formation of the IDRs (intrinsically disordered regions) contained in MSI1. Indeed, we observed granule aggregates containing MSI1-C proteins (by α-Msi1-C) in H9 hESCs, after 48 hours of 5i naïve induction, but not in the H9 cells under mTeSR culture (Figure S4C). Interestingly, formation of granules in the hESCs were mainly related to MSI1-C proteins, as the N-terminal RRM domains (RRM1, and RRM1+RRM2) seem to fail to participate in the formation of these granules during naïve induction (Figure S4D).

**Fig. 5.**
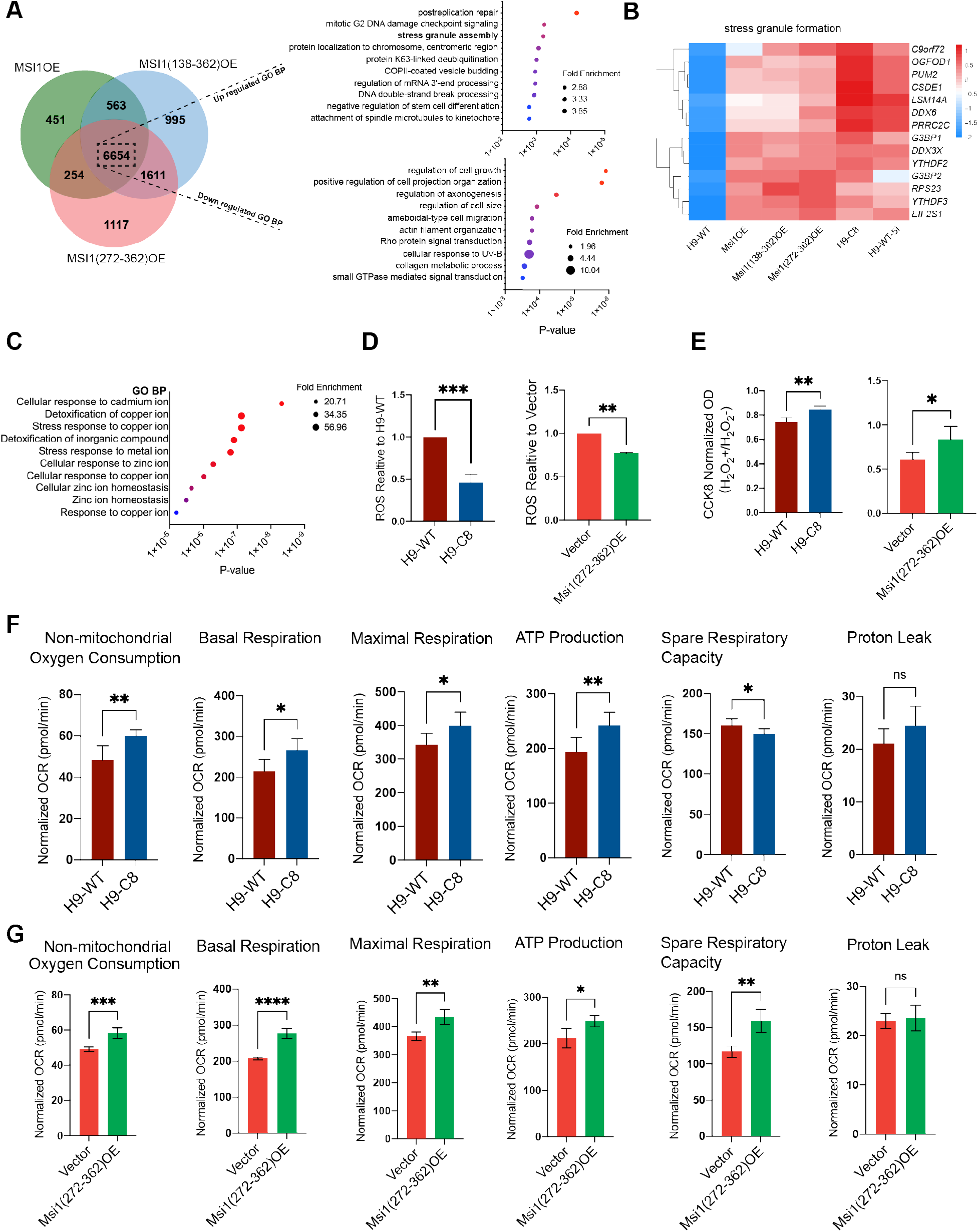
MSI1-C variants enhance stress response of hESCs with increased mitochondrial activity. (**A**) Venn diagram analysis of DEGs after overexpression of MSI1 and MSI1-C variants in H9 cells. GO analysis of the top ten biological processes for up- and down-regulated genes. (**B**) Heat map plot of DEGs involved in stress granule formation. (**C**) GO analysis of the top ten upregulated biological processes for DEGs between H9-WT Versus H9-C8. (**D**) Quantitation of ROS contents in H9-C8 relative to H9-WT cells and MSI1(272-362) overexpression in H9 relative to Vector control, calibrated by cellular protein contents. (**E**) CCK8 assay for changes in cell viability in H9-WT and H9-C8 hESCs, overexpression Vector and MSI1(272-362) in H9 hESCs after H_2_O_2_ treatment, OD values were calibrated over cellular protein contents. (**F**,**G**) Seahorse analysis of non-mitochondrial respiration, basal respiration, maximal respiration, ATP production and spare respiration capacity in H9-WT and H9-C8 cells (F), and H9 hESCs with overexpression of empty vector, Msi1, Msi1^272-362^ (G), analyzed with Mito Stress protocol. **\***indicate *P*-values (*, p<0.05; **, p<0.01; ***, p<0.001). ns, no significant difference. *t*-test, from 3 independent replicates (**F**,**G**).

Among the genes enriched in H9-C8 cells were metallothionein family genes involved in cellular response to various metals (Figure 5C), which chelate metal ions through sulfhydryl groups and scavenge oxygen radicals (Ruttkay-Nedecky *et al*, 2013). Consistently, when we detected the level of ROS, we found that H9-C8 cells, or H9 cells with MSI1^272-362^ overexpression, have lower level of ROS (Figure 5D). When treated with hydrogen peroxide (under mTeSR culture), H9-C8 cells (with endogenously expressed MSI1^272-362^) and MSI1^272-362^ overexpressed H9 cells have higher levels of cell viability, indicated by CCK8 assay (Figure 5E).

Our RIP-seq data of H9 hESCs suggested that MSI1 mainly binds to genes involved in mitochondrial activity and ATP production (Supplementary File S2). Therefore, we performed the Seahorse XF Cell Mito stress tests to examine the mitochondrial activity in hESCs. The Seahorse results showed that H9-C8, and H9-Msi1C-OE cells have higher non-mitochondrial respiration, basal respiration, maximal respiration, ATP production and spare respiration capacity (Figure 5F,G). The spare respiratory capacity was significantly increased by MSI1-C proteins, while no difference in proton leakage due to MSI1 overexpression. This suggested that the MSI1-C proteins can enhance mitochondrial activity, enabling a better cellular response to the oxidative stress.

### MSI1-C proteins bind to and upregulate DNA damage repair genes

The RNA-seq analysis showed that biological process of DNA double-strand break repair was most significantly upregulated (Figure 5A), indicating that MSI1-C proteins could potentially promote genome stability of the hESCs upon naïve induction. To further analyze this, we cross examined the RNA-seq data with RIP-seq results. The ability of RNA to interact with C-terminus of MSI1 has been predicted(Wang *et al*., 2020a), but not demonstrated. By RIP-seq analysis, we captured MSI1-C bound RNAs in H9 hESCs overexpressed with Flag-MSI1, Flag-MSI1^138-362^ and Flag-MSI1^272-362^. RIP-seq analysis showed that FL-MSI1 binds to transcripts of 2356 genes, MSI1^138-362^ binds to 1029 genes, and MSI1^272-362^ binds to 599 genes (Supplementary File S2). Further analysis against the RNA-seq data identified 1274 RNAs bound to full-length MSI1, 341 MSI1^138-362^ bound RNAs, and 190 MSI1^272-362^ bound RNAs (Supplementary File S3). Using Venn diagrams, we then identified 82 core RNAs that are bound and regulated by MSI1 and MSI1-C variants (Figure 6A, Table S4). Heat map plot of these 82 core genes showed that MSI1-C proteins confers a gene expression profile in H9 cells toward naïve induced hESCs (Figure 6B).

**Fig. 6.**
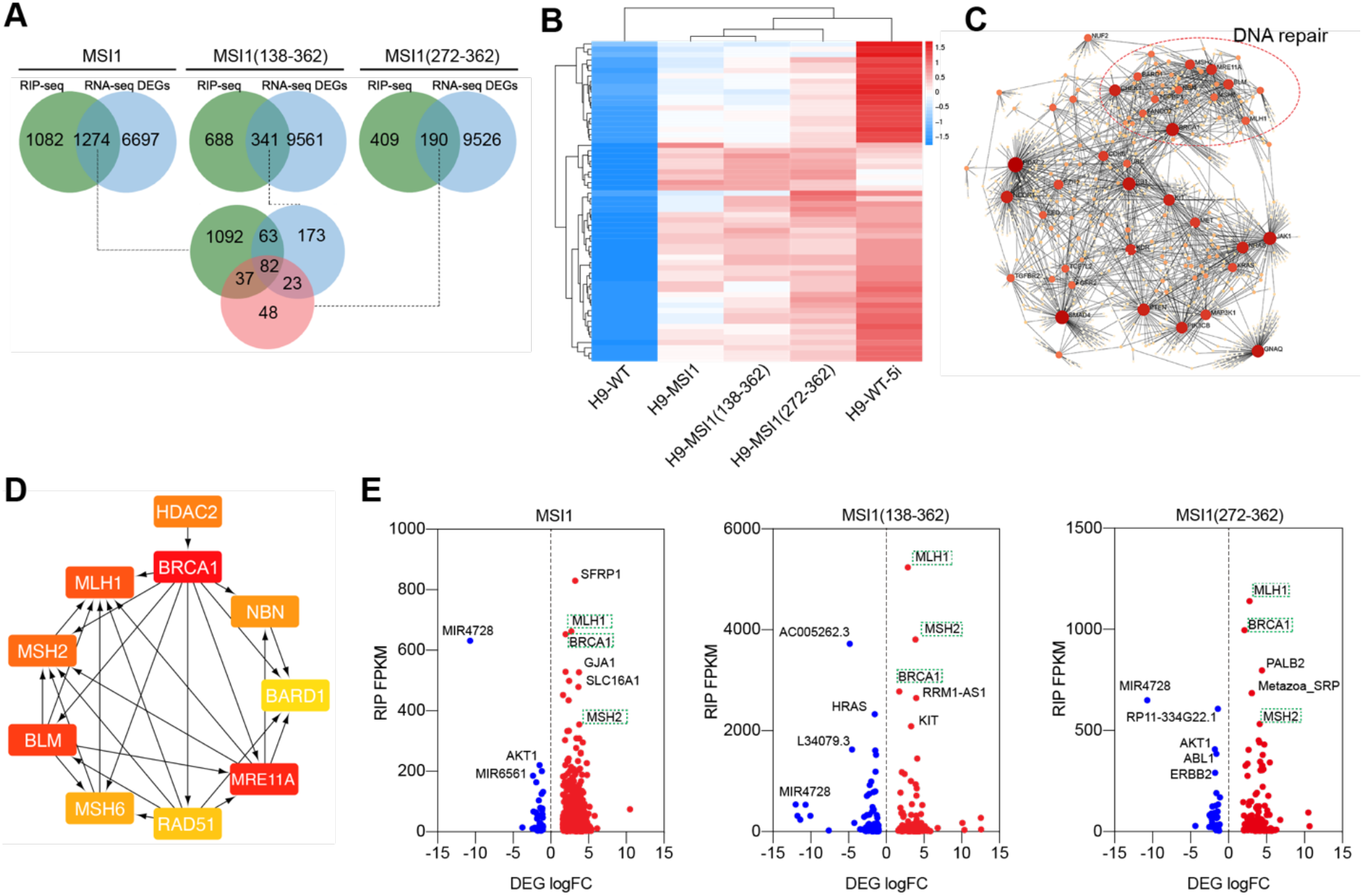
MSI1-C binds and upregulates DNA damage-repair-related genes. (**A**) Venn diagram analysis of DEGs (by RNA-seq) that are also bound by MSI1 and MSI1-C variants (by RIP-seq), in H9 hESCs overexpressing MSI1, Msi1^138-362^, Msi1^272-^ _362_. 82 core DEGs were identified. (**B**) Heatmap plot for the 82 core DEGs in H9 hESCs overexpressing MSI1, Msi1^138-362^, Msi1^272-362^ and compared to H9-WT-5i, H9-WT cells. (**C**,**D**) Protein-protein interaction (PPI) network analysis for 82 core DEGs. Each labeled point represents a core gene and the larger the node, the more interactions. The nodes in the red dashed circle are all with DNA repair function identified by GO enrichment in biology process(**C**), and top 10 hub nodes of PPI network analyzed with Cytohubba (**D**). (**E**) Dots map showing significantly upregulated (red) and downregulated (blue) genes (logFC ≤-1 or logFC ≥1, P≤0.05) in MSI1 and MSI1-C overexpressing H9 cells, against FPKM value from RIP-seq. Green dotted boxed highlight the DNA repair repair-related genes *MLH1, BRCA1* and *MSH2*, bound and regulated by MSI1-C.

We used protein association networks (Xia *et al*, 2015; Zhou *et al*, 2019) to further analyze the interactions of these 82 core proteins. In addition to signaling pathway components, such as TGFBR2, SMAD4, FGFR2, we found a prominent cluster of key protein nodes that have important roles in DNA damage repair (Figure 6C). The top 10 key nodal proteins identified are BRCA1, MLH1, MSH2, BLM, MSH6, RAD51, MRE11A, BARD1, HDAC2, and NBN (Figure 6D). Notably, all the key node proteins, except HDAC2, are in the DNA damage repair set. Plotting of the fold changes of DEGs (from RNA-seq) against the FKPM values (of RIP-seq reads) further identified the DNA repair-related genes *MLH1*(Prolla *et al*, 1994), *MSH2*(Fishel *et al*, 1994) and *BRCA1*(Zhao *et al*, 2017) as significantly up-regulated targets, that were highly bound to by both full-length MSI1 and MSI1-C proteins (Figure 6E, green dotted boxes).

### MSI1-C proteins confer formative-like pluripotency state to hESCs

The downregulation of germ cell development related genes in R1-C5 (Figure S2F) implicated a role of MSI1 in also regulating formative state transition from naïve state, as the defining characteristic of formative pluripotency is the capacity of germ stem cell differentiation. Recent studies also showed that formative PSCs bear promise in translational applications due to the capacity in interspecies chimerism formation. Thus, to put the MSI1 in the context of full range of pluripotency states, we compared our RNA-Seq data of primed state H9-WT, H9 hESCs with full-length or MSI1-C variants overexpression (in mTeSR medium), H9-C8, H9-WT-5i and H9-C8-5i cells (early stage of induction), against published results of d6 embryo (Xiang *et al*, 2020), formative PSCs (Kinoshita *et al*., 2021), and naïve induced hESCs (Khan *et al*, 2021). Principal component analysis (PCA) showed that hESCs with MSI1 or MSI1-C overexpression, even cultured in mTeSR medium, have a pluripotency state already deviated from the primed state toward the formative state (Figure 7A). Analysis of markers for pluripotency states also clustered these H9 cells with MSI1 and MSI1-C overexpression in the group of formative PSCs (Figure 7B, red bracket). This suggested that MSI1-C proteins have a “priming” effect on the transition of primed pluripotency toward naïve pluripotency.

**Fig. 7.**
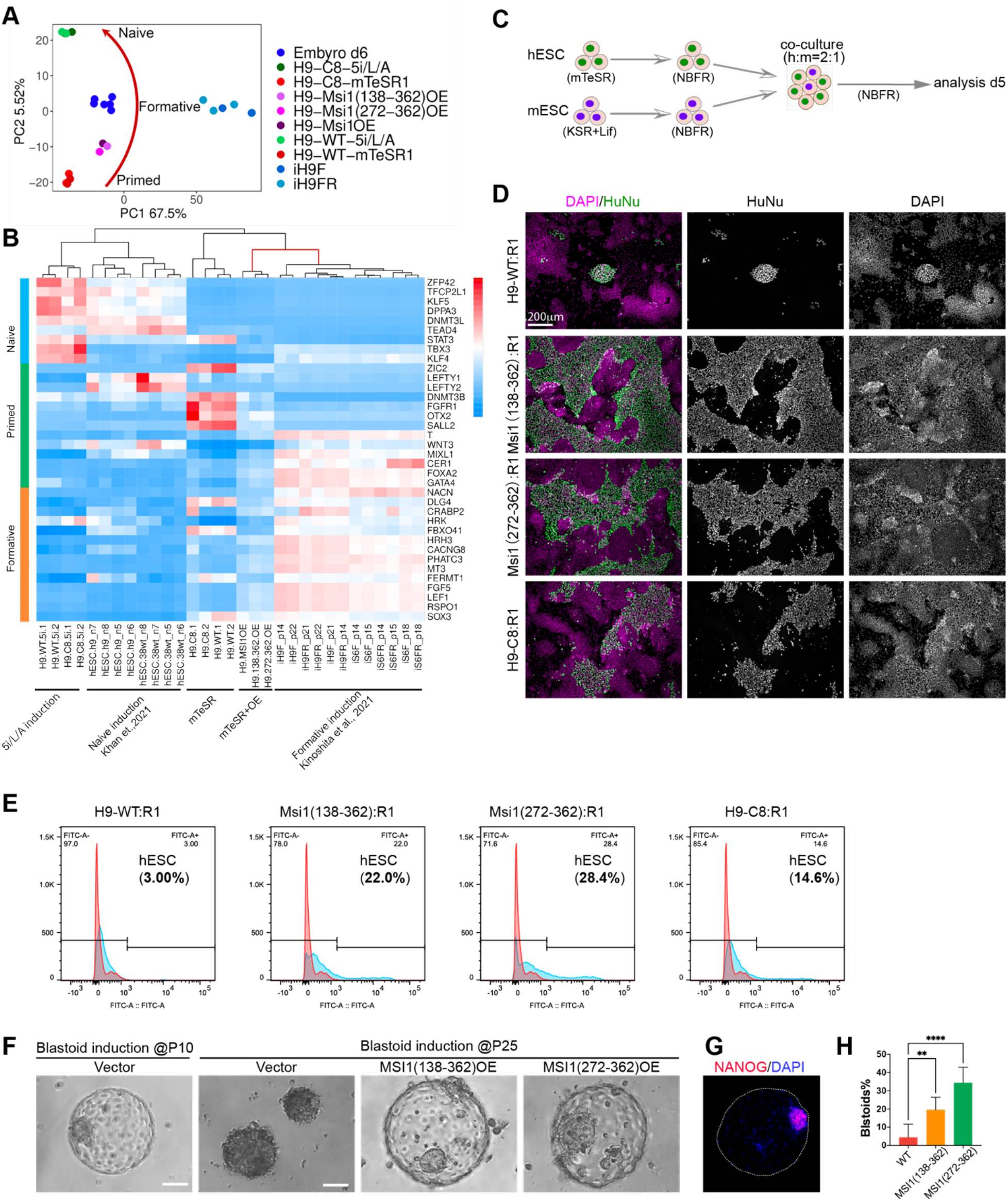
Enhanced interspecies cell competition and blastoid induction of hESCs with MSI1-C overexpression. (**A**) Principal component analysis of H9-WT, H9-C8, and H9 hESCs with overexpression of MSI1, Msi1^138-362^, Msi1^272-362^, against formative state (Embryo d6, iH9F and iH9FR) and naïve state (H9-WT-5i and H9-C8-5i) cells. (**B**) Clustering analysis of pluripotency state marker genes among H9 cells with overexpression of MSI1 and MSI1-C variants, against published data. Note H9 hESCs with overexpression of MSI1, MSI1-C variants are clustered together with formative PSCs (red bracket). (**C**,**D**) Diagram and representative images of interspecies cell competition analysis. H9 and R1 cells were acclimated to NBFR medium before co-culture at ratio of 2:1 (**C**). H9 cells detected with anti-Human nuclear antigen (HuNu) were shown in green, nuclei stained with DAPI were shown in purple (**D**). Scale bar, 200 μm. (**E**) Flow cytometry analysis of percentage of H9 cells in the co-culture, at day 5, H9 cells detected by immunofluorescence staining of Human nuclear antigen. (**F**) Phase contrast micrographs of blastoids with vector overexpression at p10 (left), and blastoids with vector control, MSI1^138-362,^ MSI1^272-362^ overexpression at p25 (right), Scale bar 50 μm. (**G**) Example of an induced blastoid (from MSI1^272-362^ group) with immunofluorescence detection of NANOG, in the inner cell mass. (H)Percentage of successful induction of blastoids formation at p25. * indicate P-values (**, p<0.01; ****, p<0.001). t-test, from 6 independent replicates.

### MSI1-C proteins enable survival of hESCs in inter-species cell competition and increase blastoid formation capacity of induced naïve hESCs

We wondered whether the upregulated expression of DNA damage repair genes (Figure 6), enhanced stress response and mitochondrial activity (Figure 5) by MSI1-C proteins have any positive effect on the survival and proliferation of hESCs. We first addressed the question with an inter-species cell competition assay. We cultured the H9 and R1 cells on Matrigel at a ratio of 2:1, and analyzed the percentage of H9 cells, at 1 day and 5 days of co-culture (Figure 7C). Both H9 and R1 cells were transitioned to the NBFR medium, which allows a suitable state for both cell types. The R1 colonies became flatten and tight, with typical morphology of primed stem cells (Figure S5A). As cell competition is contact dependent (Zheng *et al*, 2021), we seeded the H9 and R1 cells in a higher density so that at 1 day after plating, the H9 and R1 cells were at direct contact, with comparable percentages of H9 cells in the co-culture (Figure S5B).

Consistent to the previous report (Zheng *et al*., 2021), in the day 5 co-culture few H9-WT hESCs survived, with occasional H9 colonies in the space not occupied by the R1 cells (Figure 7D, Figure S5C). However, there were plenty of H9 cells when Msi1-C was overexpressed, with some R1 cells found inside the H9 colonies (Figure 7D). Flow cytometry analysis confirmed that Msi1^138-362^ and Msi1^272-362^ overexpression increased the percentage of H9 cells to 22.0% and 28.4%, from 3.0% (H9-WT), respectively (Figure 7E). H9-C8 cells, which have endogenous MSI1-C proteins, also showed higher competition capacity, compared to H9-WT cells (14.6% vs 3.0%, Figure 7D,E). This demonstrated that MSI1-C proteins could enhance the survival and proliferation of hESCs in interspecies cell competition.

It was reported that prolonged suppression of the MAPK pathway via the MEK1/2 inhibitor (PD0325901 in 2i/LIF for mouse stem cell or 5i/L/A for human stem cell) lead to chromosomal abnormalities, and impairs the developmental potential of naïve stem cell(Choi *et al*, 2017; Di Stefano *et al*, 2018). As MSI1 and MSI1-C bound and down-regulated the MAPK antagonist mir4728(Schmitt *et al*, 2015) and up-regulated DNA repair genes MLH1, MSH2 and BRAC1 (Figure 6E), which may reduce MEK1/2 inhibition effect and preserves genomic stability in naïve induced H9 cells. Thus, we used blastoid induction assay as a model to examine whether the hESCs with MSI1-C expression have higher ability to form induced blastoids (Yanagida *et al*, 2021). All the cells were induced and cultured in the 5i/L/A system, for a prolonged period. We maintained the H9 hESCs with MSI1-C overexpression for over 25 passages (to date). Under 5i/L/A culture condition, the H9-WT cells at passage 25 could barely form any blastoid (4.4%). But both H9-Msi1(272-362)-OE and H9-Msi1(272-362)-OE cells could still form blastoids (19.5% and 34.4%, respectively), with typical morphology of induced blastoid, and expression of NANOG in the inner cell mass (Kagawa *et al*, 2022)(Figure 7F-H). Under 5i naïve induction system, it has been shown that the success rate of blastoid induction of naïve induced PSCs decreases significantly after 20 passages (Yu *et al*., 2021a). Thus, this observation manifested that the MSI1-C proteins can increase the survival and proliferation of hESCs, leading to enhanced performance in blastoid induction.

In summary, this study demonstrates that Msi1 have important functions in regulating the pluripotency states in ESCs. There exist MSI1-C proteins in naïve R1 mESCs, but not in primed state H9-WT hESCs. Both MSI1 and MSI1-C proteins regulate pluripotency states in ESCs, and the MSI1-C proteins facilitate primed-to-naïve pluripotency transition. MSI1-C proteins confers better cellular response to environmental stresses and higher expression of DNA repair genes, enables better survival and proliferation of hESCs that can be induced to form blastoid.

## DISCUSSION

### MSI1 as double swords in pluripotency regulation

The RNA binding protein MSI1 was named after the heroic Japanese samurai *Miyamoto Musashi* (Nakamura *et al*, 1994), who was known to fight with two swords, a long sword, the *katana*, and a short sword, the *wakizashi*. MSI1 is often abnormally activated in tumor cells and is considered a target for cancer therapeutics (Glazer *et al*, 2008; Wang *et al*, 2010; Wang *et al*, 2013). On the other hand, MSI1 regulates asymmetric division of progenitor cells in the homeostasis of the nervous system and the intestine (Chiremba & Neufeld, 2021; Kaneko *et al*, 2000; Kanemura *et al*, 2001; Maria Cambuli *et al*, 2013), and its upregulation is often associated with tissue repair and organ regeneration (Kinoshita *et al*, 2019; Wang *et al*, 2020b). Thus, MSI1 has multifaceted functions. But its action in embryonic pluripotent stem cells remains elusive.

We serendipitously found that there exist short forms of MSI1 proteins that have not been reported. We identified a transcript encoding MSI1^138-362^ in R1 mESCs, and a new transcript encoding MSI1^272-362^ activated in H9 hESCs upon naïve induction (Figures 1 and 3). Like the long MSI1, these short MSI1-C proteins can bind to RNAs (Figure 6, Supplementary File S3) and regulate pluripotency. Notably, the MSI1-C proteins mainly act in naïve state ESCs, including the R1 mESCs and hESCs induced by the 5i method (Figures 1,3 and Figure S3). When overexpressed, MSI1 and MSI1-C variants promote the primed H9 hESCs toward a formative-like pluripotency state, increase hESCs’ “wining” of interspecies cell competition, maintain long-term culture of naïve induced hESCs and increase efficiency of blastoid induction (Figure 7). This is related, mechanistically, to enhanced stress resistance to environmental challenges (Figure 5). This is in consistence with MSI1 in increasing heat stress resistance (ErLin *et al*, 2015), radiation resistance (Sheng *et al*, 2020), and chemotherapy resistance(de Araujo *et al*, 2016; Lang *et al*, 2017) in cancer biology. Consequently, this should lead to more stable genomes in naïve ESCs, as indicated by the binding and elevated expression of genes involved in DNA damage repair (Figure 6). The result of higher levels of DNA repair gene expression in the naïve induced hESCs is manifested by the observation that the Msi1-C overexpressing hESCs have higher capacity to form induced blastoid (Figure 7). Thus, MSI1 regulates pluripotency states in ESCs, both through the full-length and the short C-terminus MSI1-C proteins.

### Subcellular localization of MSI1 and MSI1-C variants in PSCs

Our work shows that MSI1 and MSI1-C proteins are in the nucleus in the naïve mESCs, but the full length MSI1 is found in the cytoplasm in primed state ESCs (wild type H9). Both MSI1 and MSI1-C are present in the cytoplasm of hESCs under initial naïve induction by the 5i culture system (Figure 3). MSI1-C proteins act in the cytoplasm to regulate cellular stress resistance. Once the naïve state is established, however, MSI1-C proteins act in the nucleus to stabilize the genome (Figure 6). This altered intracellular localization of MSI1 has also been reported during inner ear hair cell homeostasis and retina development (Kinoshita *et al*., 2019; Nickerson *et al*, 2011). This highlights the importance of the subcellular localization of MSI1 and MSI1-C proteins.

The MSI1 protein has two nuclear localization sequences (NLS), in the RRMs region(Kawahara *et al*, 2011). It is possible that the NLSs are masked in the full-length MSI1 in primed state ESCs, while in naïve state, the NLS sites are available for nuclear transportation. Further analysis of the mechanism involved in regulation of the subcellular localization, and the regulation of MSI1-C expression, should provide more insights for the action of MSI1 in PSCs.

### Implication for MSI1 in embryonic development and interspecies chimerism

As an RNP, MSI1 binds to RNA target through two RRMs(de Araujo *et al*., 2016; Lin *et al*, 2019), and the N-terminal RRM1 is more important than RRM2 in binding MSI1 targets(Miyanoiri *et al*, 2003). Thus, when researchers investigated the role of MSI1 in the mouse central nervous system, the RRM1 was the major target (Sakakibara *et al*., 2002).The C-terminal sequences of MSI1 has received little attention. This is because like other RNPs, the MSI1-C terminus contains IDRs (intrinsically disordered regions), the structure of which is still difficult to determine. Thus, our understanding of the action of MSI1, so far, has been benefited mainly from studying the RRMs of MSI1, or the full-length MSI1, like the long sword *Miyamoto Musashi* used. In lieu of our discovery of the MSI1-C proteins in ESCs, it will necessitate using additional loss-of-function models to specifically target the MSI1-C variants, to fully understand the function of Mushashi-1 in embryonic development. This is an interesting direction that we are further investigating.

Further understanding of MSI1 in embryonic development may also pave the way for utilizing MSI1-C variant for better chimerism formation. The higher level of survival of hESCs in our interspecies cell competition assay suggests that the MSI1^138-362^ and MSI1^272-362^ variants, in addition to others such as *P65*^*KO*^ and *MYD88* ^*KO*^ (Zheng *et al*., 2021), maybe reasonable candidates for optimizing chimeric embryos derivation.

## Acknowledgements

Research in the authors’ lab was supported by the National Key R&D Program of China (2021YFA0805000, 2020YFA0803604) and National Natural Science Foundation of China (31571491, 31771608, and 31970778).

## Author Contributions

Ying Chen, Youwei Chen and Gufa Lin conceived the study. Youwei Chen performed cell culture, animal procedures, molecular cloning, immunofluorescence, Western blot, RIP, RIP-seq experiment, collected and analyzed data; Hailin Zhang, Jiazhen Han generated DNA constructs and performed Western blot analysis. Qianyan Li performed cell cultures. Ying Chen and Gufa Lin supervised the study. Youwei Chen, Ying Chen and Gufa Lin analyzed data and wrote the manuscript.

## Declaration of Interest

The authors declare no competing interests.

## METHODS

### Cell cultures

H9 hESCs were cultured using mTeSR medium (Stemcell technologies, Cambridge, USA). Cells were cultured on Matrigel (Corning)-coated culture dishes, with daily medium change. H9 hESCs were passaged as small cluster using Versene solution (Thermo Fisher Scientific, USA). R1 mESCs were cultured on mouse embryonic fibroblast feeder cells inactivated with mitomycin-C (Sigma-Aldrich), using mouse ESC (mESC) medium (knockout DMEM, Life Technologies, Carlsbad, CA, USA) supplemented with 15% knockout serum replacer (Life Technologies, Carlsbad, CA, USA), 1×GlutaMAX (Life Technologies, Carlsbad, CA, USA), 1×Anti-Anti (Life Technologies, Carlsbad, CA, USA), 1% MEM non-essential amino acids (Life Technologies, Carlsbad, CA, USA), 0.1 mM 2-Mercaptoethanol (Sigma-Aldrich, St. Louis, MO, USA) and 1000 Units/ml ESGRO-mLIF (Millipore, Burlington, MA, USA), at 37°C in 5% CO2. R1 mESCs were passaged as single cell using 0.05%-trypsin-EDTA (Life Technologies, Carlsbad, CA, USA), for R1-C5 passaging, dissociated single cells were place in cell culture dish incubated for 45 minutes at 37°C, 5% CO_2_ to allow the differentiated cells to adhere (depleted), and then the suspended cells were collected for passaging.

### Primed pluripotency state induction of R1 mESC

Induction of R1 mESC to primed state was performed following the protocol reported previously (Zhang et al., 2010). Briefly, R1 cells were dissociated into single cells using 0.05% trypsin-EDTA (Life Technologies, Carlsbad, CA, USA), resuspended in MEF culture medium (DMEM supplemented with 10% fetal bovine serum (FBS, Thermo Fisher, USA), 1% 1×GlutaMAX, 1×Anti-Anti, 1% MEM non-essential amino acids), and deprived off feeder cells by incubation at 37°C for 45 minutes. The feeder cell-deprived R1 cells were resuspended in MES medium without LIF to form suspension aggregates, which were dissociated two days later with 0.05% trypsin-EDTA and plated on culture dishes coated with FBS. Cells were cultured in DMEM/F12 containing10% KSR, 20 ng/ml activin A (R&D Systems) and 12 ng/ml bFGF (R&D Systems). Flattened colonies were manually picked for passaging.

### Naïve pluripotency state induction of H9 hESC

H9 cells were induced to the naïve state using the 5i method (Theunissen et al., 2014). Briefly, H9 cells maintained on Matrigel were seeded onto feeder cells for two generations for acclimation with mTeSR medium, and were dissociated into single cells using Accutase (Thermo Fisher, USA). Dissociated H9 cells (2 × 10^5^, for one well of a 12-well plate) were cultured with mTeSR supplemented with 10 μM Y27632 (Stemcell technologies, Cambridge, USA) for one day before switching to the 5i medium (DMEM/F12 : Neural basal = 1:1 mix, 1×N, 1× B27, 1×GlutaMAX, 1% NEAA, 0.1 mM BME, 1 μM PD0325901, 3 μM Chir99021, 0.5 μM SB590885, 1 μM WH-4-023, 10 µm Y27632, 20 ng/mL hLIF, 20 ng/mL Activin A, 8 ng/mL FGF2). Induced naïve state colonies with translucent edges were visible after 10 days of culture, and passaged using Accutase (Thermo Fisher, USA).

### Blastoid induction

Blastoid induction was performed as previously described (Yanagida *et al*., 2021). In brief, H9 hESCs cultured with 5i medium were dissociated with Acctuase, pelleted in washing medium (DMEM/F12 supplemented with 0.1% BSA), and resuspended in PD+A83+Y medium (N2B27 supplemented with 1.5 μM PD0325901, 1 μM A83-01 and 10 μM Y-27632). Then 200 cells were dispensed into each well of an ultra-low attachment 96-well plate (Corning Coster). The plates were centrifuged at 300 g for 4 min at room temperature to cluster cells at the bottom of the wells. After 48 hours, medium was replaced with pre-warmed N2B27 supplemented with 0.5 μM A83-01. Blastoid forms after 18-24 hours of medium replacement.

### Cell competition assay

Cell competition assay was performed as reported previously (Zheng et al., 2021). Briefly, H9 cells cultured medium were switched from mTeSR1 to NBFR medium (DMEM/F12 and Neurobasal medium mixed at 1:1 ratio, 0.5 × N2, 0.5× B27, 2 mM GlutaMax, 1× nonessential amino acids, 0.1 mM 2-mercaptoethanol, 20 ng/ml FGF2, 2.5 μM IWR1, and 1 mg/ml BSA) and passaged 3 times for adaption. Mouse R1 cell were cultured on feeders in NBFR medium and passaged onto plates coated with Matrigel after feeder cell depletion. For cell competition assay, primed H9 and R1 cells were dissociated into single cells using Accutase and seeded at a 2:1 ratio at a density of 5×10^4^ per well of 24-well plate, which was pre-coated with Matrigel. Cells were fixed with 4% PFA at day 1 and day 5 for Immunofluorescence analysis.

### MSI1 deletion using CRISPR-Cas9 system

Synthesized gRNAs prepared with Precision gRNA Synthesis Kit (Thermo Fisher) were mixed with Cas9 protein (Thermo Fisher) to form a complex, and electroporated into cells using the Neon transfection system (Thermo Fisher) using the following parameters: 1200 V/20 ms/2 plus for H9; 1600 V/10 ms/3 plus for R1. Single cell was seeded into 96-well plate using limited dilution method. A portion of the formed clone was used to extract genomic DNA for PCR analysis. Deletions were confirmed by sequencing of the PCR products.

### siRNA transfection

siRNA sequences are listed in Supplementary Table S5. Lipofectamine® 3000 (Thermo Fisher) Reagent was used for siRNA transfection. In brief, hESCs were dissociated into single cell by Accutase (Thermo Fisher), 1×10^5^ cells were seeded into 1 well of 12-well plate pre-coated with Matrigel and cultured in mTeSR with 10μM Y27632. Next day, medium was replaced with TeSR1 without Y27632, and 30 pmol of siRNA mixed with 3μL of Lipofectamine 3000 was added to the cells.

### MSI1 and MSI1-C overexpression with Lentivirus

For overexpression of MSI1 and MSI1-C, lentivirus was prepared from 293FT cells cultured with DMEM high glucose containing 10% FBS, 1%NEAA, 1% GlutaMax and 1% Anti-Anti. Cells with 70%~80% confluence were transfected with 10 μg MSI1-FL or MSI1-C (MSI1^138-362^, MSI1^272-362^), 6 μg psPAX2 and 4 μg pMD2.G plasmids (psPAX2 and pMD2.G were gift from Didier Trono, Addgene plasmid # 12260) by X-tremeGENE™ HP DNA Transfection Reagent (Roche). At 36- and 56-hours post-transfection, supernatants were collected and centrifuged (16000 g, 4°C, 1hour) after adding 40% PEG6000 and 4M NaCl to final concentration of 8%~10% PEG6000 and 0.3M NaCl (Marino et al., 2003). Virus pellets were aliquoted and stored at −80°C. For lentiviral transfection, (1-2) × 10^5^ cells dissociated by Accutase (H9) or Trypsin-EDTA (R1 were mixed with 50-200 μL of lentiviral particles for 12 hours, in suspension. Cells were seeded onto Matrigel coated plate (H9), or feeder-cell plated culture plate (R1). Efficiency of transfection was observed 48 hours later.

### Immunofluorescence staining

Immunofluorescence staining was performed as previously described (Chen et al., 2017). In brief, cells were fixed in 4% PFA (Sigma-Aldrich) and permeabilized with 1% Triton X-100 (Sigma-Aldrich) in PBS for 10 minutes, and blocked with blocking buffer (PBS containing 1% BSA and 0.3% Triton X-100). After incubation of primary antibody, cells were washed and incubated with secondary antibodies (1:500-1000 dilution), washed with PBS containing 0.1% Tween-20, and counterstained with DAPI. Immunofluorescence data were analyzed by Zenblue (Zeiss), or captured with DMI-6000B DM-250 microscopes equipped with digital cameras (Leica). Antibodies used are listed in the Key Resource Table (supplementary Table S6).

### Western Blotting

Cells were collected and homogenized with RIPA buffer (Thermo Scientific) containing complete protease inhibitors (Roche). For nuclear and cytoplasmic fractionation, the Qproteome Nuclear Protein Kit (QIAGEN) was used. The protein concentration was quantified by BAC protein assay kit (Thermo Scientific). Cell Lysate was denatured by adding 6 × sample buffer containing 330 mM Tris-Cl pH 6.8, 30% glycerol,1 mM EDTA, 9 % SDS, 550 mM DTT, 0.3% bromophenol blue and heated for 10 minutes at 90°C. Proteins were separated on SDS-polyacrylamide gels and processed for conventional western blotting. The primary and secondary antibodies (listed in Key Resource Table) were diluted in TBST (Tris: 20 mM, NaCl: 150 mM, Tween® 20 detergent: 0.1% w/v) containing 5% (w/v) skimmed milk. Peroxidase activity was detected by chemiluminescence (Tanon, High-sig ECL Western Blotting Substrate, Cat#180-501) and captured with Amersham Imager 600 (General Electric). Antibodies used are listed in the Key Resource Table (supplementary Table S6).

### RT-PCR, RT-qPCR and 5’RACE

Total RNA was extracted with Trizol reagent (Life Technologies), treated with Turbo DNase I (Ambion), and reverse-transcribed into complementary DNA with the Superscript IV RT system (Life Technologies). GoTaq® G2 Master Mixes (Promega) were used for RT-PCR and Fuji was used for grayscale quantification of RT-PCR product bands on Agarose gel. RT-PCR primers are listed in Supplementary table S5.

For RT-qPCR, TransScript® Uni All-in-One First-Strand cDNA Synthesis SuperMix for qPCR kit (TransGen, AU341) were used for cDNA synthesis. SYBR Green PCR Master Mix was used for RT-qPCR and detection in a Lightcycler (Roche). For detection of the short full-length and short MSI1 transcripts, 5’ RACE was performed as previously reported (Scotto-Lavino et al., 2006). cDNAs were synthesized by the Superscript IV RT system with GSP-RT primer and purified by QIAquick spin columns (Qiagen). Ploy-A was added by terminal deoxynucleotidyl transferase (TdT, Invitrogen) supplement with dATP. Ploy-A cDNA was then used for nest-PCR amplification of the MSI1 transcripts. The 5’ RACE products were separated using gel electrophoresis, purified and subjected to TA cloning and DNA sequencing.

### Mito stress assay

The mitochondrial stress test is performed using the Seahorse XF Cell Mito Stress Test Kit (Agilent, 103015-100), according to the manufacturer’s instructions, as described previously (Zhang *et al*, 2018). In brief, cells were seeded onto 96-well Seahorse plates pre-coated with Matrigel at density of 40 × 10^4^ cells per well. Culture media were switched to Seahorse XF DMEM medium (supplemented with 1 mmol/L sodium pyruvate, 2 mmol/L-glutamine and 10 mmol/L) 1 hour before assay. Selective inhibitors were injected during the measurements to achieve final concentrations of oligomycin (2.5 μM), FCCP (300 nM), antimycin (2.5 μM), rotenone (2.5 μM). The OCR values were normalized to the Protein quantity in each well, quantified by BAC protein assay kit (Thermo Scientific, Cat#23227). Changes in OCR response to the addition of inhibitors were defined as the maximal change after the chemical injection compared with the last OCR value before the injection.

### ROS analysis

Quantitative ROS assays were conducted with OxiSelect In Vitro ROS/RNS Assay Kit (Cell Biolabs, Inc.; San Diego, California) according to the manufacturer’s instructions. Briefly, cells were collected and homogenized in PBS (final concentration is 30 mg/mL) and centrifuged (10,000 g, 5 min, 4°C). Samples and standards were assayed in triplicates on 96-well plates. 50 μL of samples or standards were added into the assay plate wells and mixed with 50 μL of Catalyst. After a 5-minute incubation at room temperature, 100 μL of DCFH solution was added to each well and the plate was kept in dark for 40 minutes before the fluorescence was measured with a plate reader at 480 nm excitation / 530 nm emission. ROS content was calculated based on the standard curve. Cells were collected from 3 batches of cultures independently.

### RNA immunoprecipitation

RNA immunoprecipitation was performed with reversible cross-linking of the cells, as previously described (Niranjanakumari *et al*, 2002). In brief, 1×10^7^ cells were washed three times with pre-chilled PBS and resuspended in 10 mL of PBS, cells were fixed with 1% formaldehyde (diluted from 37% HCHO/10% methanol stock solution, Sigma-Aldrich) at room temperature for 10 minutes with slow mixing. Crosslinking reactions was quenched by the addition of glycine (pH 7.0) to a final concentration of 0.25 M, followed by incubation at room temperature for 5 minutes. The cells were harvested by centrifugation and washed twice with ice-chilled PBS. Fixed cells were resuspended in 2 mL of RIPA buffer containing protease inhibitors (Roche) and RNAase inhibitors (Ambion), and sonication was used to solubilize RNPs. Insoluble material was removed by centrifugation at 16,000g for 10 minutes at 4 °C. The supernatant was mixed with the antibody and incubated overnight at 4°C with rotation, followed by incubation with protein A/G beads. Normal IgG antibodies were used as controls. The beads were collected by centrifugation at 6000 rpm for 30s and washed five times with wash buffer (50 mM Tris–Cl, pH 7.5, 1% NP-40, 1% sodium deoxycholate, 0.1% sodium dodecyl sulfate (SDS), 1 mM EDTA, 1 M NaCl and 0.2 mM PMSF). The beads containing the immunoprecipitated samples are collected and resuspended in resuspended buffer (50 mM Tris–Cl, pH 7.0, 5 mM EDTA, 10 mM DTT and 1% SDS). Samples (resuspended beads) are incubated at 70°C for 45 minutes to reverse the crosslinks. The RNA was extracted from these samples using Trizol reagent (Invitrogen) and used for RIP-PCR or RIP-Seq.

### RNA-sequencing and analysis

For transcriptome sequencing, total RNA was extracted from the cells using TRIzol® Reagent (Invitrogen) and genomic DNA was removed using DNase I (TaKara). For RIP-sequencing, RNA was extracted from RNA precipitation preparations. Then RNA quality was determined using 2100 Bioanalyser (Agilent) and quantified using the ND-2000 (NanoDrop Technologies). High-quality RNA sample (OD260/280=1.8~2.2, OD260/230 ≥2.0, RIN≥6.5, 28S:18S≥1.0, >10μg) is used to construct sequencing library. RNA-seq transcriptome libraries were prepared following TruSeq™ RNA sample preparation Kit from Illumina (San Diego, CA), using 1 μg of total RNA. Messenger RNA was isolated with poly-A selection by oligo(dT) beads and fragmented using fragmentation buffer. cDNA synthesis, end repair, A-base addition and ligation of the Illumina-indexed adaptors were performed according to Illumina’s protocol. Libraries were then size selected for cDNA target fragments of 200–300 bp on 2% Low Range Ultra Agarose (Sigma-Aldrich) followed by PCR amplified using Phusion DNA polymerase (NEB) for 15 PCR cycles. After quantification with TBS380, paired-end libraries were sequenced by Illumina NovaSeq 6000 sequencing (150bp*2, Shanghai BIOZERON Co., Ltd). The raw paired end reads were trimmed and quality controlled by Trimmomatic with parameters (SLIDINGWINDOW:4:15 MINLEN:75). Then clean reads were separately aligned to reference genome with orientation mode using hisat2 (https://ccb.jhu.edu/software/hisat2/index.shtml) software. This software was used to map with default parameters. The quality assessment of these data was done by qualimap_v2.2.1(http://qualimap.bioinfo.cipf.es/). htseq (https://htseq.readthedocs.io/en/release_0.11.1/) was used to count each gene reads. To identify DEGs (differential expression genes) between the two different samples, the expression level for each gene was calculated using the fragments per kilobase of exon per million mapped reads (FRKM) method. R statistical package edgeR (Empirical analysis of Digital Gene Expression in R) was used for differential expression analysis. The DEGs between two samples were selected using the following criteria: the logarithmic of fold change was greater than 2 and the false discovery rate (FDR) should be less than 0.05. Heatmaps were generated using the R ‘pheatmap’’ function. Panther was applied for GO enrichment analysis. Fisher’s exact test was used for enrichment tests.

### PCA plot

Formative stage and embryo D6 genes counts were obtained from GSE131551 and GSE136447 (Kinoshita *et al*., 2021; Xiang *et al*., 2020). Zero count genes were omitted. Normalization, and differential expression analyses were performed using the R/Bioconductor edgeR package. Normalized counts were transformed into fragments per million (FPKM). Principal component analyses (PCA) were performed using the R ‘prcomp’ function based on log2-transformed Z-score expression values.

## QUANTIFICATION AND STATISTICAL ANALYSIS

Cells were observed under a Leica DM16FC or DMI6000 microscope. Images were taken with digital cameras (Leica) and Figures were prepared with Adobe Photoshop (Adobe). Unpaired *t*-tests were used for comparison of means of data, between groups. Data were presented as mean +/-standard deviation. Differences were considered significant if P-value <0.05(*). P-values were presented with asterisks in Figures; <0.05(*), <0.01(**), <0.001(***), or <0.001(****),.

## DATA AND SOFTWARE AVAILABILITY

The datasets reported in this paper can be accessed at https://www.ncbi.nlm.nih.gov/geo/query/acc.cgi?acc=GSE197608 The GEO accession number is GSE197608.

## Notes

### Competing Interest Statement

The authors have declared no competing interest.

## REFERENCES

Cambuli FM, Correa BR, Rezza A, Burns SC, Qiao M, Uren PJ, Kress E, Boussouar A, Galante PA, Penalva LO et al (2015) A Mouse Model of Targeted Musashi1 Expression in Whole Intestinal Epithelium Suggests Regulatory Roles in Cell Cycle and Stemness. Stem Cells 33: 3621–3634

Chan YS, Goke J, Ng JH, Lu X, Gonzales KA, Tan CP, Tng WQ, Hong ZZ, Lim YS, Ng HH (2013) Induction of a human pluripotent state with distinct regulatory circuitry that resembles preimplantation epiblast. Cell Stem Cell 13: 663–675

Chen TC, Huang JR (2020) Musashi-1: An Example of How Polyalanine Tracts Contribute to Self-Association in the Intrinsically Disordered Regions of RNA-Binding Proteins. Int J Mol Sci 21

Chiou GY, Yang TW, Huang CC, Tang CY, Yen JY, Tsai MC, Chen HY, Fadhilah N, Lin CC, Jong YJ (2017) Musashi-1 promotes a cancer stem cell lineage and chemoresistance in colorectal cancer cells. Sci Rep 7: 2172

Chiremba TT, Neufeld KL (2021) Constitutive Musashi1 expression impairs mouse postnatal development and intestinal homeostasis. Mol Biol Cell 32: 28–44

Choi J, Huebner AJ, Clement K, Walsh RM, Savol A, Lin K, Gu H, Di Stefano B, Brumbaugh J, Kim SY et al (2017) Prolonged Mek1/2 suppression impairs the developmental potential of embryonic stem cells. Nature 548: 219–223

de Araujo PR, Gorthi A, da Silva AE, Tonapi SS, Vo DT, Burns SC, Qiao M, Uren PJ, Yuan ZM, Bishop AJ et al (2016) Musashi1 Impacts Radio-Resistance in Glioblastoma by Controlling DNA-Protein Kinase Catalytic Subunit. Am J Pathol 186: 2271–2278

De Los Angeles A, Wu J (2021) New concepts for generating interspecies chimeras using human pluripotent stem cells. Protein Cell

Di Stefano B, Ueda M, Sabri S, Brumbaugh J, Huebner AJ, Sahakyan A, Clement K, Clowers KJ, Erickson AR, Shioda K et al (2018) Reduced MEK inhibition preserves genomic stability in naive human embryonic stem cells. Nat Methods 15: 732–740

Eggan K, Rode A, Jentsch I, Samuel C, Hennek T, Tintrup H, Zevnik B, Erwin J, Loring J, Jackson-Grusby L et al (2002) Male and female mice derived from the same embryonic stem cell clone by tetraploid embryo complementation. Nat Biotechnol 20: 455–459

ErLin S, WenJie W, LiNing W, BingXin L, MingDe L, Yan S, RuiFa H (2015) Musashi-1 maintains blood-testis barrier structure during spermatogenesis and regulates stress granule formation upon heat stress. Mol Biol Cell 26: 1947–1956

Fishel R, Ewel A, Lee S, Lescoe MK, Griffith J (1994) Binding of mismatched microsatellite DNA sequences by the human MSH2 protein. Science 266: 1403–1405

Fox RG, Park FD, Koechlein CS, Kritzik M, Reya T (2015) Musashi signaling in stem cells and cancer. Annu Rev Cell Dev Biol 31: 249–267

Gafni O, Weinberger L, Mansour AA, Manor YS, Chomsky E, Ben-Yosef D, Kalma Y, Viukov S, Maza I, Zviran A et al (2013) Derivation of novel human ground state naive pluripotent stem cells. Nature 504: 282–286

Glazer RI, Wang XY, Yuan H, Yin Y (2008) Musashi1: a stem cell marker no longer in search of a function. Cell Cycle 7: 2635–2639

Good P, Yoda A, Sakakibara S, Yamamoto A, Imai T, Sawa H, Ikeuchi T, Tsuji S, Satoh H, Okano H (1998) The human Musashi homolog 1 (MSI1) gene encoding the homologue of Musashi/Nrp-1, a neural RNA-binding protein putatively expressed in CNS stem cells and neural progenitor cells. Genomics 52: 382–384

Huang K, Zhu Y, Ma Y, Zhao B, Fan N, Li Y, Song H, Chu S, Ouyang Z, Zhang Q et al (2018) BMI1 enables interspecies chimerism with human pluripotent stem cells. Nat Commun 9: 4649

Kagawa H, Javali A, Khoei HH, Sommer TM, Sestini G, Novatchkova M, Scholte Op Reimer Y, Castel G, Bruneau A, Maenhoudt N et al (2022) Human blastoids model blastocyst development and implantation. Nature 601: 600–605

Kaneko Y, Sakakibara S, Imai T, Suzuki A, Nakamura Y, Sawamoto K, Ogawa Y, Toyama Y, Miyata T, Okano H (2000) Musashi1: an evolutionally conserved marker for CNS progenitor cells including neural stem cells. Dev Neurosci 22: 139–153

Kanemura Y, Mori K, Sakakibara S, Fujikawa H, Hayashi H, Nakano A, Matsumoto T, Tamura K, Imai T, Ohnishi T et al (2001) Musashi1, an evolutionarily conserved neural RNA-binding protein, is a versatile marker of human glioma cells in determining their cellular origin, malignancy, and proliferative activity. Differentiation 68: 141–152

Kawahara H, Okada Y, Imai T, Iwanami A, Mischel PS, Okano H (2011) Musashi1 cooperates in abnormal cell lineage protein 28 (Lin28)-mediated let-7 family microRNA biogenesis in early neural differentiation. J Biol Chem 286: 16121–16130

Khan SA, Park KM, Fischer LA, Dong C, Lungjangwa T, Jimenez M, Casalena D, Chew B, Dietmann S, Auld DS et al (2021) Probing the signaling requirements for naive human pluripotency by high-throughput chemical screening. Cell Rep 35: 109233

Kilens S, Meistermann D, Moreno D, Chariau C, Gaignerie A, Reignier A, Lelievre Y, Casanova M, Vallot C, Nedellec S et al (2018) Parallel derivation of isogenic human primed and naive induced pluripotent stem cells. Nat Commun 9: 360

Kinoshita M, Barber M, Mansfield W, Cui Y, Spindlow D, Stirparo GG, Dietmann S, Nichols J, Smith A (2021) Capture of Mouse and Human Stem Cells with Features of Formative Pluripotency. Cell Stem Cell 28: 453–471 e458

Kinoshita M, Fujimoto C, Iwasaki S, Kashio A, Kikkawa YS, Kondo K, Okano H, Yamasoba T (2019) Alteration of Musashi1 Intra-cellular Distribution During Regeneration Following Gentamicin-Induced Hair Cell Loss in the Guinea Pig Crista Ampullaris. Front Cell Neurosci 13: 481

Lang Y, Kong X, He C, Wang F, Liu B, Zhang S, Ning J, Zhu K, Xu S (2017) Musashi1 Promotes Non-Small Cell Lung Carcinoma Malignancy and Chemoresistance via Activating the Akt Signaling Pathway. Cell Physiol Biochem 44: 455–466

Lee JH, Laronde S, Collins TJ, Shapovalova Z, Tanasijevic B, McNicol JD, Fiebig-Comyn A, Benoit YD, Lee JB, Mitchell RR et al (2017) Lineage-Specific Differentiation Is Influenced by State of Human Pluripotency. Cell Rep 19: 20–35

Lin JC, Tsai JT, Chao TY, Ma HI, Liu WH (2019) Musashi-1 Enhances Glioblastoma Migration by Promoting ICAM1 Translation. Neoplasia 21: 459–468

Maria Cambuli F, Rezza A, Nadjar J, Plateroti M (2013) Brief report: musashi1-eGFP mice, a new tool for differential isolation of the intestinal stem cell populations. Stem Cells 31: 2273–2278

Miyanoiri Y, Kobayashi H, Imai T, Watanabe M, Nagata T, Uesugi S, Okano H, Katahira M (2003) Origin of higher affinity to RNA of the N-terminal RNA-binding domain than that of the C-terminal one of a mouse neural protein, musashi1, as revealed by comparison of their structures, modes of interaction, surface electrostatic potentials, and backbone dynamics. J Biol Chem 278: 41309–41315

Nakamura M, Okano H, Blendy JA, Montell C (1994) Musashi, a neural RNA-binding protein required for Drosophila adult external sensory organ development. Neuron 13: 67–81

Nickerson PE, Myers T, Clarke DB, Chow RL (2011) Changes in Musashi-1 subcellular localization correlate with cell cycle exit during postnatal retinal development. Exp Eye Res 92: 344–352

Niranjanakumari S, Lasda E, Brazas R, Garcia-Blanco MA (2002) Reversible cross-linking combined with immunoprecipitation to study RNA-protein interactions in vivo. Methods 26: 182–190

Ohyama T, Nagata T, Tsuda K, Kobayashi N, Imai T, Okano H, Yamazaki T, Katahira M (2012) Structure of Musashi1 in a complex with target RNA: the role of aromatic stacking interactions. Nucleic Acids Res 40: 3218–3231

Prolla TA, Pang Q, Alani E, Kolodner RD, Liskay RM (1994) MLH1, PMS1, and MSH2 interactions during the initiation of DNA mismatch repair in yeast. Science 265: 1091–1093

Ruttkay-Nedecky B, Nejdl L, Gumulec J, Zitka O, Masarik M, Eckschlager T, Stiborova M, Adam V, Kizek R (2013) The role of metallothionein in oxidative stress. Int J Mol Sci 14: 6044–6066

Sakakibara S, Imai T, Hamaguchi K, Okabe M, Aruga J, Nakajima K, Yasutomi D, Nagata T, Kurihara Y, Uesugi S et al (1996) Mouse-Musashi-1, a neural RNA-binding protein highly enriched in the mammalian CNS stem cell. Dev Biol 176: 230–242

Sakakibara S, Nakamura Y, Yoshida T, Shibata S, Koike M, Takano H, Ueda S, Uchiyama Y, Noda T, Okano H (2002) RNA-binding protein Musashi family: roles for CNS stem cells and a subpopulation of ependymal cells revealed by targeted disruption and antisense ablation. Proc Natl Acad Sci U S A 99: 15194–15199

Schmitt DC, Madeira da Silva L, Zhang W, Liu Z, Arora R, Lim S, Schuler AM, McClellan S, Andrews JF, Kahn AG et al (2015) ErbB2-intronic microRNA-4728: a novel tumor suppressor and antagonist of oncogenic MAPK signaling. Cell Death Dis 6: e1742

Sheng X, Lin Z, Lv C, Shao C, Bi X, Deng M, Xu J, Guerrero-Juarez CF, Li M, Wu X et al (2020) Cycling Stem Cells Are Radioresistant and Regenerate the Intestine. Cell Rep 32: 107952

Shibata S, Umei M, Kawahara H, Yano M, Makino S, Okano H (2012) Characterization of the RNA-binding protein Musashi1 in zebrafish. Brain Res 1462: 162–173

Smith A (2017) Formative pluripotency: the executive phase in a developmental continuum. Development 144: 365–373

Susaki K, Kaneko J, Yamano Y, Nakamura K, Inami W, Yoshikawa T, Ozawa Y, Shibata S, Matsuzaki O, Okano H et al (2009) Musashi-1, an RNA-binding protein, is indispensable for survival of photoreceptors. Experimental Eye Research 88: 347–355

Theunissen TW, Powell BE, Wang H, Mitalipova M, Faddah DA, Reddy J, Fan ZP, Maetzel D, Ganz K, Shi L et al (2014) Systematic identification of culture conditions for induction and maintenance of naive human pluripotency. Cell Stem Cell 15: 471–487

Wang J, Lan L, Wu X, Xu L, Miao Y (2020a) Mechanism of RNA recognition by a Musashi RNA-binding protein. bioRxiv: 2020.2010.2030.362756

Wang Q, Lin Y, Sheng X, Xu J, Hou X, Li Y, Zhang H, Guo H, Yu Z, Ren F (2020b) Arachidonic Acid Promotes Intestinal Regeneration by Activating WNT Signaling. Stem Cell Reports 15: 374–388

Wang XY, Penalva LO, Yuan H, Linnoila RI, Lu J, Okano H, Glazer RI (2010) Musashi1 regulates breast tumor cell proliferation and is a prognostic indicator of poor survival. Mol Cancer 9: 221

Wang XY, Yu H, Linnoila RI, Li L, Li D, Mo B, Okano H, Penalva LO, Glazer RI (2013) Musashi1 as a potential therapeutic target and diagnostic marker for lung cancer. Oncotarget 4: 739–750

Xia J, Gill EE, Hancock RE (2015) NetworkAnalyst for statistical, visual and network-based meta-analysis of gene expression data. Nat Protoc 10: 823–844

Xiang L, Yin Y, Zheng Y, Ma Y, Li Y, Zhao Z, Guo J, Ai Z, Niu Y, Duan K et al (2020) A developmental landscape of 3D-cultured human pre-gastrulation embryos. Nature 577: 537–542

Yanagida A, Spindlow D, Nichols J, Dattani A, Smith A, Guo G (2021) Naive stem cell blastocyst model captures human embryo lineage segregation. Cell Stem Cell 28: 1016–1022 e1014

Yoda A, Sawa H, Okano H (2000) MSI-1, a neural RNA-binding protein, is involved in male mating behaviour in Caenorhabditis elegans. Genes Cells 5: 885–895

Yu L, Wei Y, Duan J, Schmitz DA, Sakurai M, Wang L, Wang K, Zhao S, Hon GC, Wu J (2021a) Blastocyst-like structures generated from human pluripotent stem cells. Nature 591: 620–626

Yu L, Wei Y, Sun HX, Mahdi AK, Pinzon Arteaga CA, Sakurai M, Schmitz DA, Zheng C, Ballard ED, Li J et al (2021b) Derivation of Intermediate Pluripotent Stem Cells Amenable to Primordial Germ Cell Specification. Cell Stem Cell 28: 550–567 e512

Zhang K, Li L, Huang C, Shen C, Tan F, Xia C, Liu P, Rossant J, Jing N (2010) Distinct functions of BMP4 during different stages of mouse ES cell neural commitment. Development 137: 2095–2105

Zhang M, Chen Y, Xu H, Yang L, Yuan F, Li L, Xu Y, Chen Y, Zhang C, Lin G (2018) Melanorctin receptor 4 (Mc4r) regulates limb regeneration. Dev Cell 46: 397–409. e395

Zhao W, Steinfeld JB, Liang F, Chen X, Maranon DG, Jian Ma C, Kwon Y, Rao T, Wang W, Sheng C et al (2017) BRCA1-BARD1 promotes RAD51-mediated homologous DNA pairing. Nature 550: 360–365

Zhao XY, Li W, Lv Z, Liu L, Tong M, Hai T, Hao J, Guo CL, Ma QW, Wang L et al (2009) iPS cells produce viable mice through tetraploid complementation. Nature 461: 86–90

Zheng C, Hu Y, Sakurai M, Pinzon-Arteaga CA, Li J, Wei Y, Okamura D, Ravaux B, Barlow HR, Yu L et al (2021) Cell competition constitutes a barrier for interspecies chimerism. Nature 592: 272–276

Zhou G, Soufan O, Ewald J, Hancock REW, Basu N, Xia J (2019) NetworkAnalyst 3.0: a visual analytics platform for comprehensive gene expression profiling and meta-analysis. Nucleic Acids Res 47: W234–W241

