## supplementary information for "Musashi1 and its short C-terminal variants regulate pluripotency states in embryonic stem cells"

**Running title**: Musashi1 regulates embryonic stem cell pluripotency

### SUPPLEMENTARY FIGURES

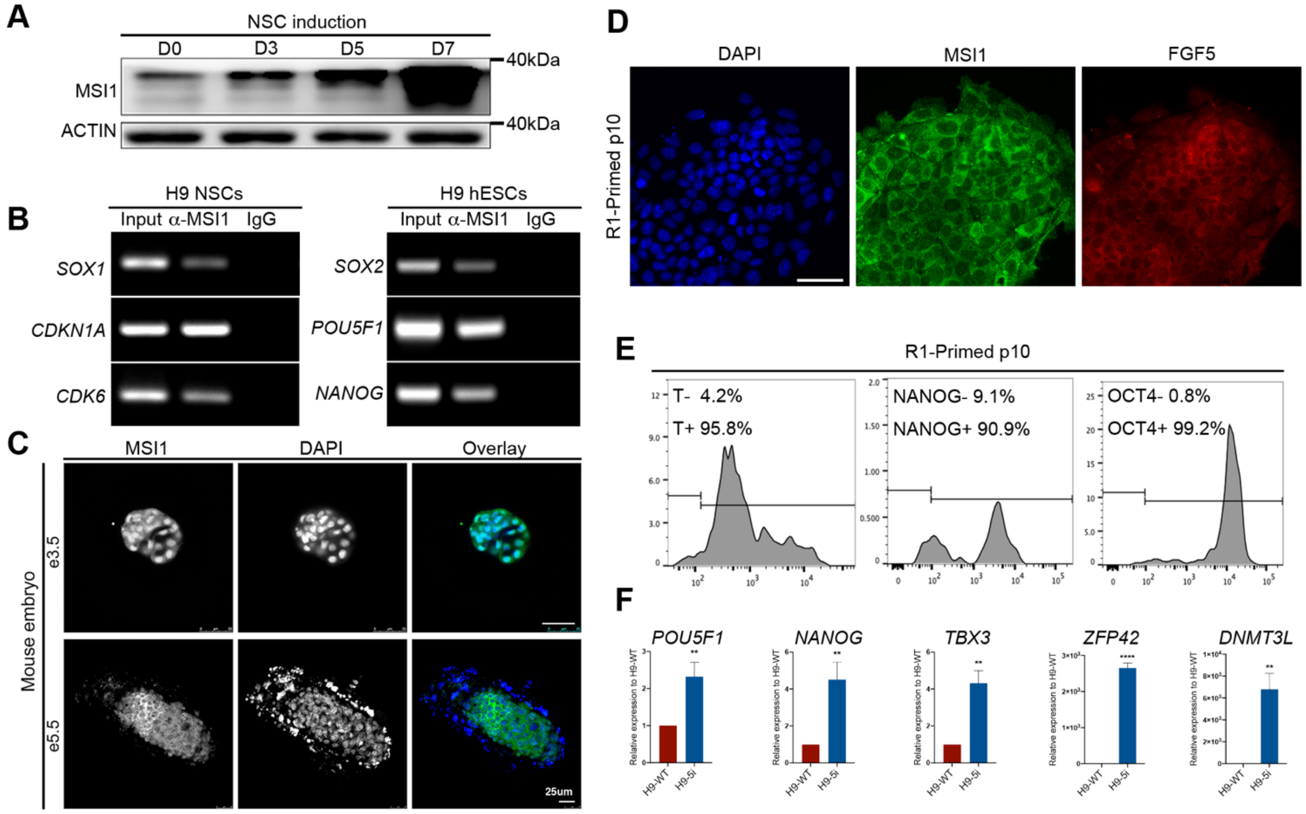


**Fig.S1 MSI1 expression in NSC, H9 hESCs, pre- and post- implantation mouse embryos, and naïve induced H9 cells, related to Figure 1**

(**A**) Western blot detection for MSI1 expression during NSC induction from H9 hESCs, from day 0 (D0), D3, D5 to D7. Note MSI1 is present at D0. ACTIN as loading control.
(**B**) RIP-PCR for MSI1 bound RNAs in NSC and H9 hESCs. MSI1 binds to *SOX1*, *CDKN1A* and *CDK6* in NSC, and binds to *SOX2*, *POU5F1*, *NANOG* in H9 hESCs.
(**C**) Immunofluorescence detection of MSI1 (α-MSI1, abcam ab52865) in pre-implantation (e3.5) and post-implantation (e5.5) mouse embryos, showing different locations of MSI1 signals. Note the mainly nuclear localized signals of MSI1 in e3.5 embryo. Scale bar, 25μm.
(**D**) Immunofluorescence detection of MSI1 (α-MSI1, abcam ab52865), FGF5 (ab88118) in R1-primed p10 cells. Scale bar, 50 μm.
(**E**) Flow cytometry detection of primed state marker T and pluripotency marker NANOG, OCT4. Histogram is shown. Gating for negative staining is based unstained cells.
(**F**) RT-qPCR detection of pluripotency genes *POU5F1*, *NANOG*, naïve state genes *TBX3*, *ZFP42*, *DNMT3L* after induction of H9 hESC to naïve state (H9-5i), compared to H9 wild type controls (H9-WT).

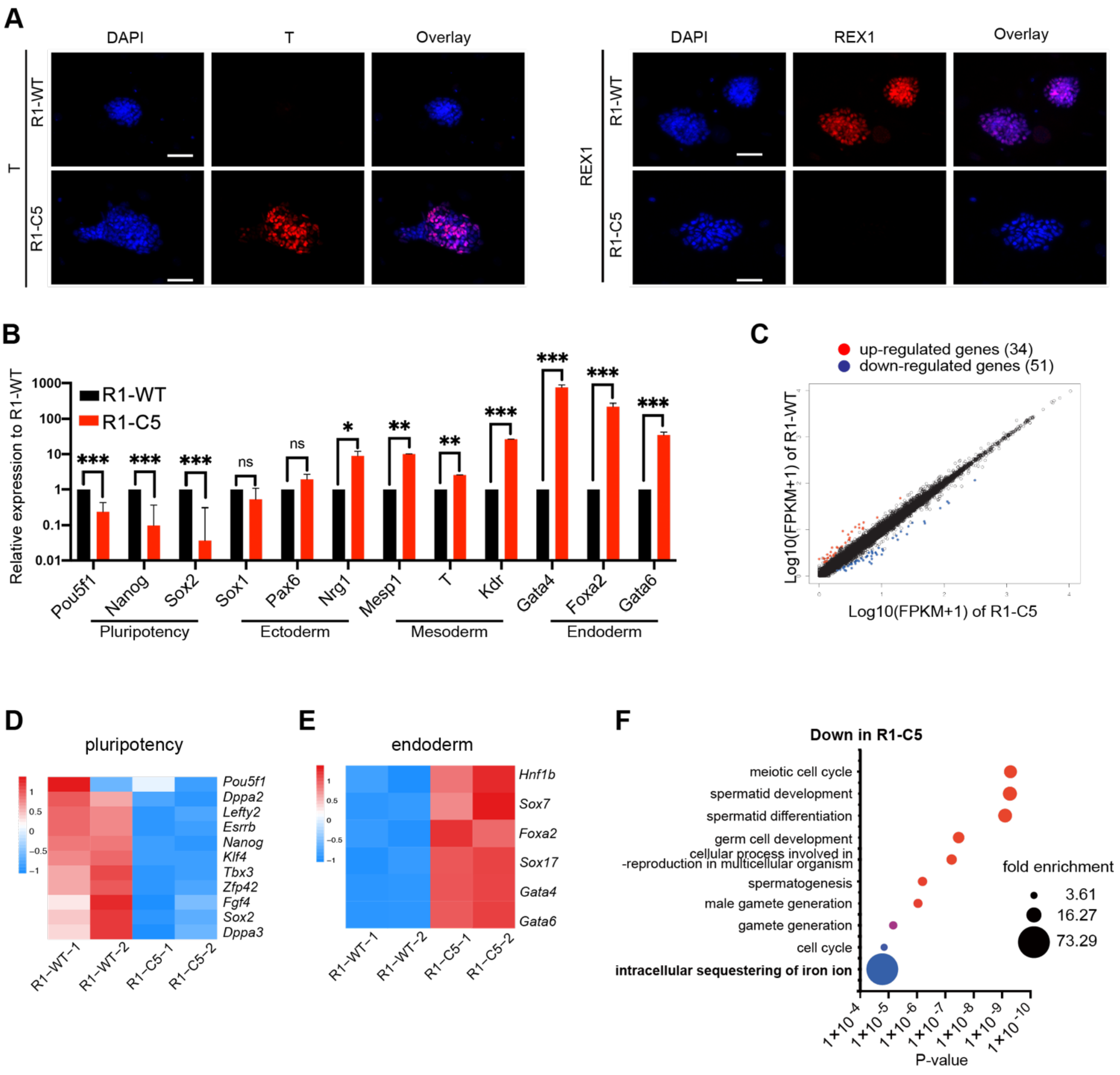


**Fig.S2 Loss-of-function of MSI1 in mESCs, related to Figure 2**

(**A**) Immunofluorescence detection of T and REX1 expression in R1-WT and R1-C5 cells. Scale bar, 50 μm.

(**B**) RT-qPCR detection of *Pou51f*, *Nanog*, *Sox2*; ectodermal genes *Sox1*, *Pax6*, *Nrg1*; mesodermal genes *Mesp1*, *T*, *Kdr* and endodermal genes *Gata4*, *Foxa2*, *Gata6* in R1-WT and R1-C5 cells. Asterisks indicate P-values. (*, p<0.05, **, p<0.01, ***, p<0.001). ns, no significant difference. t-test, from 3 independent replicates.

(**C**) Scatted plot of DEGs in R1-C5, with 34 and 51 up-, down- regulated genes.

(**D,E**) Heatmap plot of DEGs involved in pluripotency (**D**), endoderm development (**E**).

(**F**) GO enrichment analysis of biological processes, down-regulated in R1-C5 cells.

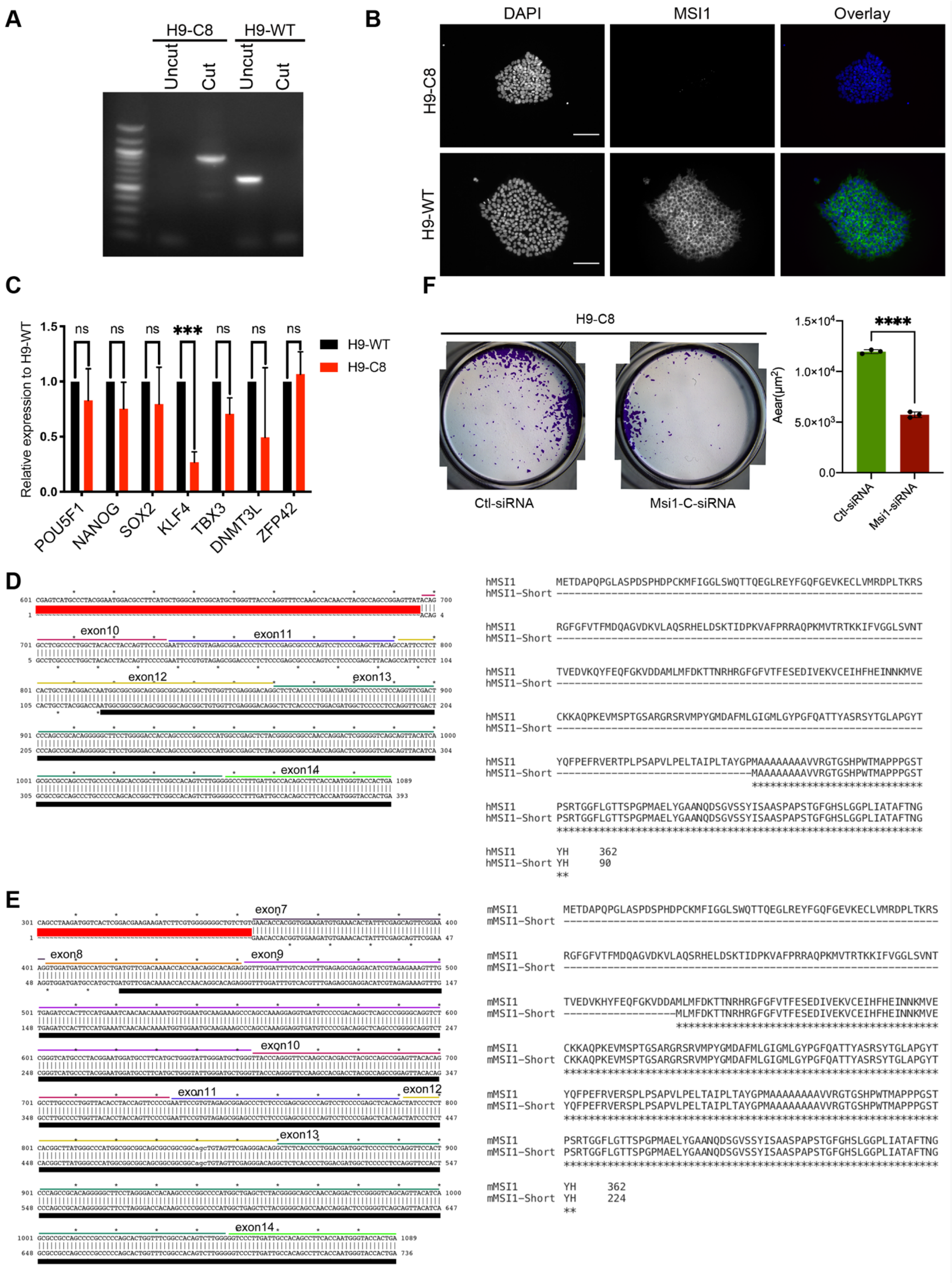
A
H9-C8
H9-WT
Uncut
Cut
Uncut
Cut
B
DAPI
MSI1
Overlay
H9-C8
H9-WT
C
Relative expression to H9-WT
1.5
1.0
0.5
0.0
ns
***
H9-WT
H9-C8
POU5F1
NANOG
SOX2
KLF4
TBX3
DNMT3L
ZFP42
F
H9-C8
Ctl-siRNA
Msi1-C-siRNA
Aear(μm²)
1.5×10⁴
1.0×10⁴
5.0×10³
0.0
****
Ctl-siRNA
Msi1-siRNA
D
exon10
exon11
exon12
exon13
exon14
hMSI1 METDAPQPGLASPDSPHDPCKMFIGGLSWQTTQEGLREYFGQFGEVKECLVMRDPLTKRS
hMSI1 RGFGFVTFMDQAGVDKVLAQSRHELDSKTIDPKVAFPRRAQPKMVTRTKKIFVGGLSVNT
hMSI1 TVEDVKQYFEQFGKVDDAMLMFDKTTNRHRGFGFVTFESEDIVEKVCEIHFHEINNKMVE
hMSI1 CKKAQPKEVMSPTGSARGRSRVMPYGMDAFMLGIGMLGYPGFQATTYASRSYTGLAPGYT
hMSI1 YQFPEFRVERTPLPSAPVLPELTAIPLTAYGPMAAAAAAAAVVRGTGSHPWTMAPPPGST
hMSI1-Short MAAAAAAAAVVRGTGSHPWTMAPPPGST
hMSI1 PSRTGGFLGTTSPGPMAELYGAANQDSGVSSYISAASPAPSTGFGHSLGGPLIATAFTNG
hMSI1-Short PSRTGGFLGTTSPGPMAELYGAANQDSGVSSYISAASPAPSTGFGHSLGGPLIATAFTNG
hMSI1 YH 362
hMSI1-Short YH 90
E
exon7
exon8
exon9
exon10
exon11
exon12
exon13
exon14
mMSI1 METDAPQPGLASPDSPHDPCKMFIGGLSWQTTQEGLREYFGQFGEVKECLVMRDPLTKRS
mMSI1 RGFGFVTFMDQAGVDKVLAQSRHELDSKTIDPKVAFPRRAQPKMVTRTKKIFVGGLSVNT
mMSI1 TVEDVKHYFEQFGKVDDAMLMFDKTTNRHRGFGFVTFESEDIVEKVCEIHFHEINNKMVE
mMSI1-Short MLMFDKTTNRHRGFGFVTFESEDIVEKVCEIHFHEINNKMVE
mMSI1 CKKAQPKEVMSPTGSARGRSRVMPYGMDAFMLGIGMLGYPGFQATTYASRSYTGLAPGYT
mMSI1-Short CKKAQPKEVMSPTGSARGRSRVMPYGMDAFMLGIGMLGYPGFQATTYASRSYTGLAPGYT
mMSI1 YQFPEFRVERSPLPSAPVLPELTAIPLTAYGPMAAAAAAAAAVVRGTGSHPWTMAPPPGST
mMSI1-Short YQFPEFRVERSPLPSAPVLPELTAIPLTAYGPMAAAAAAAAAVVRGTGSHPWTMAPPPGST
mMSI1 PSRTGGFLGTTSPGPMAELYGAANQDSGVSSYISAASPAPSTGFGHSLGGPLIATAFTNG
mMSI1-Short PSRTGGFLGTTSPGPMAELYGAANQDSGVSSYISAASPAPSTGFGHSLGGPLIATAFTNG
mMSI1 YH 362
mMSI1-Short YH 224

**Fig.S3 MSI1-C variants in ESCs, related to Figure 3**
(**A**) Genomic PCR detection of H9-C8, a clone with expected deletion of human MSI1.
(**B**) Immunofluorescence detection of MSI1 (with ab52865) show absence of MSI1 expression in H9-C8 hESCs. Scale bar, 100 μm.
(**C**) RT-qPCR detection of pluripotency genes *POU5F1*, *NANOG*, *SOX2*, *KLF4*, and naïve state genes *TBX3*, *DNMT3L*, *ZFP42* in H9-WT and H9-C8 hESCs. ***, $p<0.001$.
(**D**) Sequencing results of the small MSI1-C transcripts obtained by 5'RACE in H9-5i hESCs, aligned to human MSI1 mRNA (left). The human MSI1-C transcript has an ORF (underlined) that would encode a protein corresponding to hMSI1(272-362) (right).
(**E**) Sequencing results of the small MSI1-C transcript obtained by 5'RACE in R1 mESCs in naïve state, aligned to mouse MSI1 mRNA (left). The mouse Msi1-C transcript has an ORF that would encode a protein corresponding to mMSI1(138-362) (right).
Positions of corresponding exons on the transcripts are indicated above the sequences. ORFs are underlined with thick black lines in (**D**,**E**).

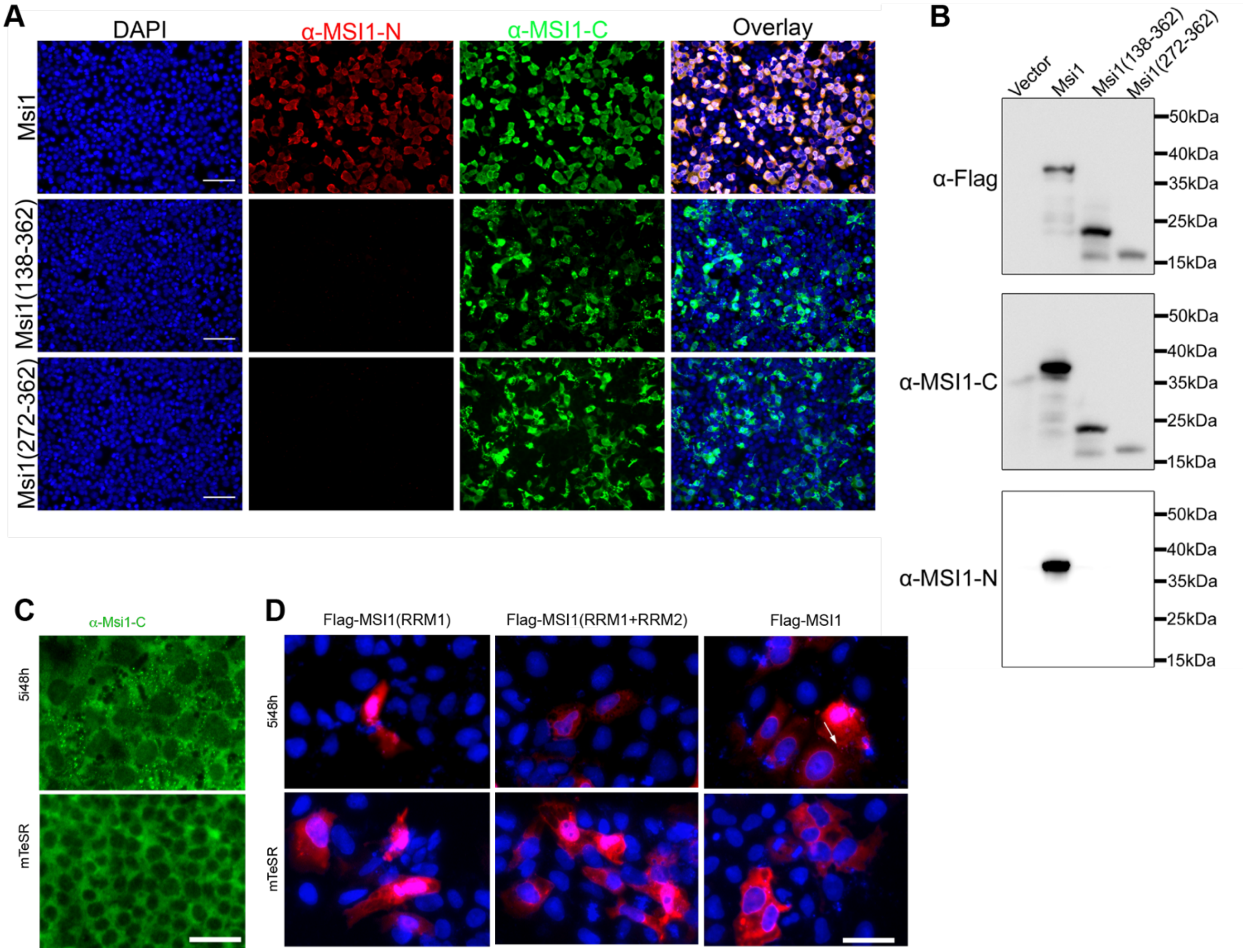


**Fig.S4 Overexpression of MSI1-C variants in hESCs, related to Figure 4**

(**A**,**B**) Immunofluorescence and Western blot detection of MSI1 and MSI1-C variants in 293FT cells with overexpression, showing the translation of MSI1-C variants, and the specificity of α-MSI1-N (ab21628), α-MSI1-C (GT2377) antibodies. Note that part of $MSI1^{138-362}$ can produce further shortened proteins. Note Flag tags were added to both the N- and C- terminus of the overexpression constructs, to detect potential variants. Scale bar, 50 µm.

(**C**) Immunofluorescence detection of MSI1 with α-MSI-C (GT2377) in H9 hESCs, in mTeSR1 or after 48h of 5i induction.

(**D**) Immunofluorescence detection of Flag-tag in H9 hESCs with overexpression of Flag-MSI1 (RRM1), Flag-MSI1 (RRM1+RRM2) and Flag-MSI1, under mTeSR culture or after 48h of 5i induction. White arrow indicates rare granules in Flag-MSI1 overexpressing hESCs.

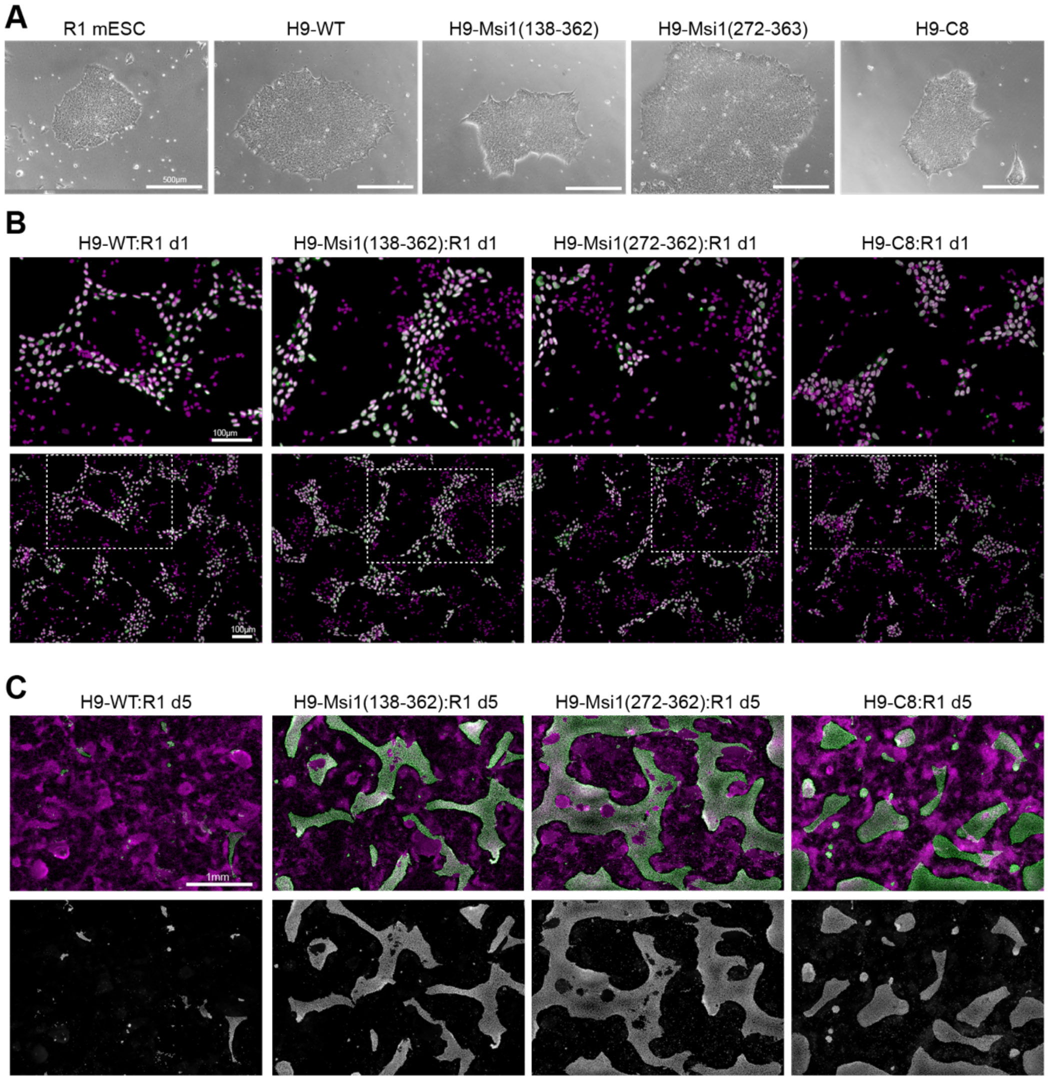


**Fig.S5 Cell competition analysis of hESCs, related to Figure 7**

(**A**) Phase contrast micrographs of R1 and H9 cells, adapted to NBFR medium for co-culture analysis.
(**B**) Representative image of co-culture of H9 and R1 cells, 1 day after seeding, at high density to allow cell contact. H9 detected by anti-HuNu shown in green, overlayed with DAPI shown in purple. Images in upper rows were higher magnification of areas outlined in the lower row.
(**C**) Representative image of co-culture of H9 and R1 cells, 5 day after seeding. Multiple images were aligned to show a wide field of the cells. Upper row: overlay images. Lower row: H9 cells detected by anti-HuNu.
Scale bar, 0.5 mm in (**A**), 100 μm in (**B**), 1mm in (**C**).

#### SUPPLEMENTARY TABLES AND FILES

##### Table S1. Summary of CRISPR/Cas9 trials of MSI1 deletion in H9 hESCs

| Experiment | Target | Exon | Number of clones picked | Number of successful knockouts | Number of passable clones |
|---|---|---|---|---|---|
| 1 | MSI1 | Exon1-Exon14 | 10 | 3 | 0 |
| | MSI1 | Exon1-Exon9 | 10 | 1 | 0 |
| | MSI1 | Exon1-Exon7 | 10 | 3 | 3 |
| 2 | MSI1 | Exon1-Exon14 | 6 | 1 | 0 |
| | MSI1 | Exon1-Exon9 | 6 | 1 | 0 |
| | MSI1 | Exon1-Exon7 | 3 | 1 | 1 |
| 3 | MSI1 | Exon1-Exon14 | 5 | 2 | 0 |

##### Supplementary File S2. RNA-seq and RIP-seq data.xlxs

| MSI1 RIP gene FPKM in H9 | MSI1 RIP gene FPKM in R1 | RNA-seq H9-C8 vs H9-WT | RNA-seq R1-C5 vs R1-WT | RNA-seq MSI1 MSI1C -OE | RIP-seq H9 MSI1 MSI1-C -OE Flag | RIP-peak H9 MSI1-FL OE | RIP-peak H9 MSI1(138-362)OE | RIP-peak H9 MSI1(272-362)OE |
|---|---|---|---|---|---|---|---|---|

Results are listed in excel file: RNA-seq and RIP-seq data.xlxs

##### Supplementary File S3. MSI1 and MSI1-C binging and regulated genes.xlsx

Results are listed in excel file: MSI1 and MSI1-C binging and regulated genes.xlsx

#### Table S4. List of 82 core genes FPKM (Figure 6A)

| Gene name | Gene id | H9-WT | H9-MSI1 OE | H9-MSI1(138-362)OE | H9-MSI1(272-362)OE |
|---|---|---|---|---|---|
| ERRFI1 | ENSG00000116285 | 0.59 | 6.456 | 5.388 | 6.731 |
| JAK1 | ENSG00000162434 | 0.092 | 1.323 | 1.482 | 1.947 |
| NRAS | ENSG00000213281 | 2.514 | 29.948 | 37.675 | 39.396 |
| RIT1 | ENSG00000143622 | 0.794 | 4.504 | 5.8 | 4.563 |
| NUF2 | ENSG00000143228 | 0.079 | 2.847 | 3.681 | 6.311 |
| LGALS8 | ENSG00000116977 | 0.169 | 3.155 | 2.969 | 3.559 |
| NPM1P24 | ENSG00000215086 | 0.075 | 2.106 | 1.349 | 1.399 |
| PTEN | ENSG00000171862 | 0.294 | 2.149 | 2.849 | 4.096 |
| TCF7L2 | ENSG00000148737 | 0.856 | 8.051 | 9.01 | 9.409 |
| FGFR2 | ENSG00000066468 | 0.965 | 3.71 | 5.244 | 5.49 |
| RPS6KA4 | ENSG00000162302 | 15.573 | 4.739 | 5.076 | 4.598 |
| EED | ENSG00000074266 | 0.314 | 6.971 | 7.066 | 5.579 |
| MRE11 | ENSG00000020922 | 0.127 | 2.056 | 2.135 | 2.879 |
| nan | ENSG00000254422 | 1.133 | 4.434 | 4.743 | 15.133 |
| CHEK1 | ENSG00000149554 | 0.631 | 6.447 | 6.11 | 7.436 |
| KRAS | ENSG00000133703 | 0.56 | 9.078 | 9.198 | 10.811 |
| RB1 | ENSG00000139687 | 0.13 | 1.704 | 1.713 | 2.743 |
| AKT1 | ENSG00000142208 | 32.429 | 11.6 | 7.167 | 9.235 |
| USP8 | ENSG00000138592 | 0.107 | 1.076 | 1.44 | 1.884 |
| BLM | ENSG00000197299 | 0.397 | 3.892 | 2.512 | 3.709 |
| TSC2 | ENSG00000103197 | 31.561 | 6.444 | 5.389 | 5.929 |
| PALB2 | ENSG00000083093 | 0.187 | 3.173 | 2.98 | 3.986 |
| nan | ENSG00000261723 | 0 | 1.439 | 1.301 | 1.463 |
| ALOX12B | ENSG00000179477 | 1.172 | 0.266 | 0.414 | 0.408 |
| NCOR1 | ENSG00000141027 | 0.219 | 1.821 | 1.934 | 2.378 |
| CDK12 | ENSG00000167258 | 0.481 | 2.497 | 2.296 | 3.012 |
| MIR4728 | ENSG00000265178 | 1.66 | 0 | 0 | 0 |
| RARA | ENSG00000131759 | 3.869 | 1.534 | 0.375 | 1.119 |
| BRCA1 | ENSG00000012048 | 0.326 | 1.27 | 1.088 | 1.397 |
| BRIP1 | ENSG00000136492 | 0.103 | 1.846 | 1.624 | 2.321 |
| SMAD4 | ENSG00000141646 | 0.647 | 8.166 | 7.882 | 6.483 |
| STK11 | ENSG00000118046 | 35.982 | 10.887 | 6.124 | 6.388 |
| AKT2 | ENSG00000105221 | 13.984 | 6.099 | 2.854 | 4.544 |
| XRCC1 | ENSG00000073050 | 17.05 | 6.883 | 3.526 | 4.772 |
| POLD1 | ENSG00000062822 | 177.724 | 44.552 | 35.007 | 37.023 |
| EPCAM | ENSG00000119888 | 8.29 | 47.803 | 53.591 | 47.275 |
| MSH2 | ENSG00000095002 | 1.471 | 20.12 | 21.259 | 25.52 |
| MSH6 | ENSG00000116062 | 0.753 | 11.732 | 12.938 | 12.736 |
| FBXO11 | ENSG00000138081 | 0.393 | 11.433 | 13.479 | 10.936 |
| FANCL | ENSG00000115392 | 0.387 | 7.285 | 9.659 | 10.075 |
| NFE2L2 | ENSG00000116044 | 0.16 | 4.874 | 6.921 | 7.571 |
| PMS1 | ENSG00000064933 | 0.343 | 2.829 | 5.584 | 7.215 |
| IDH1 | ENSG00000138413 | 6.406 | 58.573 | 62.634 | 54.404 |
| BARD1 | ENSG00000138376 | 0.158 | 2.314 | 1.923 | 3.531 |
| RTEL1 | ENSG00000258366 | 11.325 | 5.051 | 2.62 | 2.482 |
| RTEL1-TNFRSF6B | ENSG00000026036 | 7.768 | 3.5 | 1.843 | 1.743 |
| RIPK4 | ENSG00000183421 | 1.163 | 0.564 | 0.528 | 0.3 |
| FANCD2 | ENSG00000144554 | 0.252 | 2.285 | 2.357 | 2.816 |
| TGFBR2 | ENSG00000163513 | 0.203 | 1.201 | 1.153 | 1.524 |
| MLH1 | ENSG00000076242 | 1.127 | 7.496 | 8.162 | 7.612 |
| MST1R | ENSG00000164078 | 1.423 | 0.568 | 0.351 | 0.556 |
| PBRM1 | ENSG00000163939 | 0.329 | 2.157 | 2.05 | 2.763 |
| TOPBP1 | ENSG00000163781 | 0.109 | 2.95 | 2.395 | 3.506 |
| PIK3CB | ENSG00000051382 | 0.136 | 2.164 | 3.031 | 3.34 |
| ETV5 | ENSG00000244405 | 0.847 | 3.921 | 3.795 | 3.448 |
| FIP1L1 | ENSG00000145216 | 0.555 | 2.32 | 2.058 | 3.157 |
| KIT | ENSG00000157404 | 0.152 | 1.389 | 1.492 | 2.053 |
| KDR | ENSG00000128052 | 0.792 | 5.822 | 3.569 | 6.307 |
| MRPS18C | ENSG00000163319 | 0.686 | 10.253 | 15.121 | 14.502 |
| ABRAXAS1 | ENSG00000163322 | 0.366 | 6.809 | 10.036 | 10.028 |
| FBXW7 | ENSG00000109670 | 0.253 | 1.75 | 1.231 | 1.738 |
| PALLD | ENSG00000129116 | 0.396 | 2.086 | 3.176 | 3.565 |
| TERT | ENSG00000164362 | 5.02 | 2.175 | 1.014 | 1.643 |
| MAP3K1 | ENSG00000095015 | 0.107 | 1.311 | 1.824 | 1.853 |
| RAD50 | ENSG00000113522 | 0.123 | 1.192 | 1.891 | 2.687 |
| HLA-H | ENSG00000206341 | 90.07 | 33.246 | 31.609 | 15.44 |
| HLA-J | ENSG00000204622 | 5.879 | 0.432 | 1.128 | 0.281 |
| HDAC2 | ENSG00000196591 | 0.974 | 9.912 | 12.717 | 13.986 |
| GOPC | ENSG00000047932 | 0.377 | 4.383 | 4.515 | 5.648 |
| CARD11 | ENSG00000198286 | 4.262 | 2.067 | 0.768 | 1.461 |
| CDK6 | ENSG00000105810 | 0.406 | 1.875 | 2.285 | 2.58 |
| MET | ENSG00000105976 | 0.107 | 1.465 | 1.718 | 1.676 |
| EZH2 | ENSG00000106462 | 0.369 | 1.891 | 2.462 | 2.627 |
| NOS3 | ENSG00000164867 | 4.094 | 1.963 | 1.841 | 1.33 |
| PPP2R2A | ENSG00000221914 | 0.361 | 3.996 | 4.724 | 5.511 |
| GGH | ENSG00000137563 | 1.123 | 12.331 | 16.11 | 13.49 |
| NBN | ENSG00000104320 | 0.202 | 2.975 | 3.22 | 3.771 |
| RAD54B | ENSG00000197275 | 0.104 | 1.858 | 3.236 | 4.542 |
| FSBP | ENSG00000265817 | 0.14 | 1.935 | 3.9 | 5.908 |
| GNAQ | ENSG00000156052 | 0.208 | 3.769 | 4.648 | 4.86 |
| ARAF | ENSG00000078061 | 20.005 | 7.645 | 7.252 | 7.476 |
| STAG2 | ENSG00000101972 | 0.638 | 8.47 | 9.983 | 11.42 |

#### Table S5. Oligonucleotide used in the current study

| Name | Purpose | | Sequence 5'-3' |
|---|---|---|---|
| Q-mTbx3 | detection | Forward | CGGATTTACTTTGGCCTTCCC |
| | | Reverse | AGGCAGTAACGGCGATGAAT |
| Q-mKdr | detection | Forward | GGATAACCTGGCTGACCCG |
| | | Reverse | GAACCACAGAGCGACAGCTA |
| Q-mEsrrb1 | detection | Forward | CCCTCTGCCCGTTCGCT |
| | | Reverse | AATCACGCGGAAGACGAGT |
| Q-mRex1 | detection | Forward | CTGGGTACGAGTGGCAGTTT |
| | | Reverse | ATCCGCAAACACCTGCTTTT |
| Q-mKlf4 | detection | Forward | AAGGATCTCGGGCAATCTGG |
| | | Reverse | TTCCTCACGCCAACGGTTA |
| Q-mDnmt3L | detection | Forward | CATCCCTACCTACGGGTTCT |
| | | Reverse | GGCTAGAGCTGTCGGAAGTC |
| Q-mT | detection | Forward | AGGTGAGCCGTCTTTCCAC |
| | | Reverse | GGTCCACAGTATGCCATCCC |
| Q-mNanog | detection | Forward | TGCTCCGCTCCATAACTTCG |
| | | Reverse | AGGCTTGTGGGGTGCTAAAA |
| Q-mGapdh | detection | Forward | TGTGAACGGATTTGGCCGTA |
| | | Reverse | ACTGTGCCGTTGAATTTGCC |
| mMsi1-N | detection | Forward | AAGGGCTGCGCGAATACTT |
| | | Reverse | CTGTGCTCTTCGAGGAAAGG |
| mMsi1-C | detection | Forward | GCACAGAGGGTTTGGATTTGTC |
| | | Reverse | TAACCCAGCATCCCAATACCC |
| Q-mSox1 | detection | Forward | TCTCCAACTCTCAGGGCTACA |
| | | Reverse | GGCTCCGACTTGACCAGAGA |
| Q-mPax6 | detection | Forward | GGTGCTGGACAATGAAAACGTATC |
| | | Reverse | GCTTTTCGCTAGCCAGGTTG |
| Q-mNrg1 | detection | Forward | AGTCACAGCTGGAGTAATGGG |
| | | Reverse | GATTTAGGGGAGCTTGGCGT |
| Q-Mesp1 | detection | Forward | CCGCCTGCCTACCCTAGAC |
| | | Reverse | CTGAAGAGCGGAGATGAGGGA |
| Q-mFoxa2 | detection | Forward | ATGCACTCGGCTTCCAGTAT |
| | | Reverse | TCACGGAAGAGTAGCCCTCG |
| Q-mGata6 | detection | Forward | GTGCCTCGACCACTTGCTAT |
| | | Reverse | CTGATGCCCCTACCCCTGAG |
| Q-mCdkn1a | detection | Forward | ATCCAGACATTCAGAGCCACAG |
| | | Reverse | CCAGACGAAGTTGCCCTCC |
| Q-mGata4 | detection | Forward | TGTGTAGCAGGCAGAAAGCA |
| | | Reverse | GTTGCTCCAGAAATCGTGCG |
| mMsi1-Full length | detection | Forward | ATGGAGACTGACGCGCCC |
| | | Reverse | TCAGTGGTACCCATTGGTGAAGG |
| hMsi1-Full length | detection | Forward | ATGGAGACTGACGCGCCC |
| | | Reverse | TCAGTGGTACCCATTGGTGAAGG |
| Q-hRex1 | detection | Forward | CTAGGCAAACCCACCCCACT |
| | | Reverse | TTCAGCAAACACCTGCTGGAC |

| Name | Purpose | | Sequence 5‘-3’ |
|---|---|---|---|
| Q-hGAPDH | detection | Forward | TCGGAGTCAACGGATTTGGT |
| | | Reverse | TTCCCGTTCTCAGCCTTGAC |
| Q-hOCT4 | detection | Forward | AGAGTGGTGACGGAGACAGG |
| | | Reverse | AAGCGATCAAGCAGCGACTA |
| Q-hCDKN1A | detection | Forward | AGTCAGTTCCTTGTGGAGCC |
| | | Reverse | CATTAGCGCATCACAGTCGC |
| Q-hTBX3 | detection | Forward | GAGGCTAAAGAACTTTGGGATCA |
| | | Reverse | CATTTCGGGGTCGGCCTTA |
| hMsi1-N | detection | Forward | GCAGACTACGCAGGAAGGG |
| | | Reverse | CGTTCGAGTCACCATCTTGGG |
| hMsi1-C | detection | Forward | CATTCCTCTCACTGCCTACGGA |
| | | Reverse | CAAGACTGTGGCCGAAGCC |
| Q-hDnmt3L | detection | Forward | TCTCAAGCTCCGTTTCACCC |
| | | Reverse | ACTTGTCCTTACATGGGGCG |
| Q-hklf4 | detection | Forward | GATGCTCACCCCACCTTCTT |
| | | Reverse | TTTCTCACCTGTGTGGGTTCG |
| Q-hSox2 | detection | Forward | GCTGCGGGCTACTGAAAAGT |
| | | Reverse | CTCCAGCTTGGGGTCCTGAA |
| Q-hCDH | detection | Forward | CATTGAGCTGTGCAACGCC |
| | | Reverse | GGCATGACACACAACAGACTC |
| BamH1-Flag-Msi1-F | expression | Forward | AAA***GGATCC***ATGGATTACAAGGATGACGACGATAAGATGGAGACTGACGCGCCC |
| BamH1-Flag-Msi1V138-F | | Forward | AAA***GGATCC***ATGGATTACAAGGATGACGACGATAAGCTGATGTTCGACAAAACCACC |
| BamH1-Flag-Msi1V272-F | | Forward | AAA***GGATCC***ATGGATTACAAGGATGACGACGATAAGGTGGAATGTAAGAAAGCTCAGCCAA |
| EcoRV-Flag-MSI1-R | | Reverse | AAA***GATATC***TCACTTATCGTCGTCATCCTTGTAATCGTGGTACCCATTGGTGAAGGCTG |
| EcoRV-Msi1(RRM1+2) | | Reverse | AAA***GATATC***TCAACCCAGCATCCCAATACCCA |
| EcoRV-Msi1(RRM1) | | Reverse | AAA***GATATCT***TCACTGTGCTCGCCGAGGGA |
| QT | 5’RACE | Forward | CCAGTGAGCAGAGTGACGAGGACTCGAGCTCAAGCTTTTTTTTTTTTTTTTT |
| Q0 | | Forward | CCAGTGAGCAGAGTGACG |
| QI | | Forward | GAGGACTCGAGCTCAAGC |
| hMsi1-GSP-RT | | RT-primer | AAATCTGAATAAAAAACAG |
| mMSi1-GSP-RT | | RT-primer | GCCAGAATTAAAAAAATC |
| hMsi1-Nest1 | | Reverse | GGCTCACTCGTGGTCCTCA |
| mMsi1-Nest1 | | Reverse | CCTCAGTCAGCTGCAGGCT |
| h/mMSI1-Nest2 | | Reverse | TCAGTGGTACCCATTGGTGAAGG |
| Uncut-hMsi1 | knockout identification | Forward | AAGGAGTGTCTGGTGATGCG |
| | | Reverse | AGAGTTCACAGAAGCCACCG |
| Cut-hMsi1 | knockout identification | Forward | ACGGAGCGAGGGCTCTAAAT |
| | | Reverse | GTGTGTCTGCAATTCGGCAA |
| Uncut-mMsi1 | knockout identification | Forward | GAATACTTCGGCCAGTTCGGG |
| | | Reverse | ACGAGAGAGAGTTCCCGGAT |
| Cut-mMsi1 | knockout identification | Forward | ATCCGAAGCCGAGCTACCTT |
| | | Reverse | TTCTCCCTTCTTCCCCACGA |
| IVT-m/hMsi1-Exon1 | gRNA synthesis | Forward | TAATACGACTCACTATAG GGGCGCGTCAGTCTCCAT |
| | | Reverse | TTCTAGCTCTAAAAC ATGGAGACTGACGCGCCC |

| Name | Purpose | | Sequence 5'-3' |
|---|---|---|---|
| IVT-hMsi1-Exon7 | gRNA synthesis | Forward | TAATACGACTCACTATAG TCACCTCGGTGCCGGTTGG |
| | | Reverse | TTCTAGCTCTAAAAC CCAACCGGCACCGAGGTGA |
| IVT-hMsi1-Exon9 | gRNA synthesis | Forward | TAATACGACTCACTATAG ATGCTGGGCATCGGCATGCT |
| | | Reverse | TTCTAGCTCTAAAAC AGCATGCCGATGCCCAGCAT |
| IVT-mMsi1-exon9 | gRNA synthesis | Forward | TAATACGACTCACTATAG TCCCCGACAGGCTCAGCCC |
| | | Reverse | TTCTAGCTCTAAAAC GGGCTGAGCCTGTCGGGGA |
| mMsi1-N-siRNA | Knockdown | Sense | CGAAGAGCACAGCCUAAGAUG |
| | | Anti-sense | UCUUAGGCUGUGCUCUUCGAG |
| m/hMsi1-C-siRNA | Knockdown | Sense | CAGCCUUCACCAAUGGGUACC |
| | | Anti-sense | UACCCAUUGGUGAAGGCUGUG |
| m/hMsi1-C-siRNA | Knockdown | Sense | CAGCCUUCACCAAUGGGUACC |
| | | Anti-sense | UACCCAUUGGUGAAGGCUGUG |
| Ctl-siRNA | Negative control | Sense | UUCUCCGAACGUGUCACGUTT |
| | | Anti-sense | ACGUGACACGUUCGGAGAATT |

**Note**: All detection PCR primers were designed with Ta at 55°C. For the ease of subsequent cloning, adaptors containing restriction enzyme sites were added and shown in ***bold italic***. Related to STAR Methods.

#### Table S6. Key resource table

| REAGENT or RESOURCE | SOURCE | IDENTIFIER |
|---|---|---|
| **Antibodies** | | |
| Recombinant Anti-Musashi 1 / Msi1 antibody [EP1302] | abcam | Cat# ab52865; RRID: AB_881168 |
| Anti-Musashi 1 C-terminus | GeneTex | Cat# GT2377; RRID: N/A |
| Anti-Musashi 1 N-terminus | abcam | Cat# ab21628 RRID: AB_2144988 |
| Anti-Musashi 2 | abcam | Cat# ab76148 RRID: AB_1523981 |
| Anti-Histone H3 antibody | abcam | Cat# ab1791; RRID: AB_302613 |
| Anti-alpha Tubulin antibody | abcam | Cat#ab7291; RRID: AB_2241126 |
| Anti-Naong antibody | CST | Cat# 3580S; RRID: AB_2150399 |
| Anti-Actin antibody | Millipore | Cat#;MAB1501R RRID: AB_2275404 |
| Anti-TBX3 antibody | abcam | Cat#;ab99302 RRID: AB_2275404 |
| Anti-Brachyury antibody | abcam | Cat#;ab209665 RRID: AB_2275404 |
| Anti-FGF5 antibody | abcam | Cat#;ab88118 RRID: AB_2275404 |
| Anti-Rex1 antibody | abcam | Cat#;ab175429 RRID: AB_2275404 |
| Anti-Oct4 antibody | CST | Cat#;2840S RRID: AB_2275404 |
| Anti-DYKDDDDK Tag antibody | Thermo Fisher Scientific | Cat# PA1-984B, RRID: AB_347227 |
| Anti-Human Nuclear Antigen antibody | abcam | Cat# ab191181 RRID: AB_2885016 |
| Donkey anti-Sheep IgG(H+L) Cross-Adsorbed Secondary Antibody, Alexa Fluor 488 | Thermo Fisher Scientific | Cat# A11015, RRID:AB_2534082 |
| Donkey anti-Rabbit IgG (H+L) Highly Cross-Adsorbed Secondary Antibody, Alexa Fluor Plus 555 | Thermo Fisher Scientific | Cat# A32794 RRID: AB_2762834 |
| Donkey anti-Mouse IgG (H+L) Highly Cross-Adsorbed Secondary Antibody, Alexa Fluor 488 | Thermo Fisher Scientific | Cat# A21202, RRID: AB_141607 |
| Goat anti-Mouse IgG (H+L) Highly Cross-Adsorbed Secondary Antibody, Alexa Fluor 555 | Thermo Fisher Scientific | Cat#A21424; RRID: AB_2275404 |
| Goat anti-Mouse IgG (H+L) Secondary Antibody, HRP | Thermo Fisher Scientific | Cat# 31430, RRID:AB_228307 |
| Goat anti-Rabbit IgG (H+L) Secondary Antibody, HRP | Thermo Fisher Scientific | Cat# 65-6120, RRID: AB_2533967 |
| Recombinant Rabbit IgG, monoclonal [EPR25A] - Isotype Control | abcam | Cat# ab172730 RRID: RRID:AB_2687931 |
| **Bacterial and Virus Strains** | | |
| DH5-alpha | TranGen Biotech | Cat# CD501 |
| **Chemicals, Peptides, and Recombinant Proteins** | | |
| Y27632 | Selleck | Cat# S1049 |
| PD0325901 | Sigma-Aldrich | Cat# 444966 |
| Chir99021 | Sigma-Aldrich | Cat# SML1046 |

| | | |
|---|---|---|
| SB590885 | Selleck | Cat# S2220 |
| WH-4-023 | Selleck | Cat# S7565 |
| A83-01 | Selleck | Cat# S7692 |
| Gossypol | Selleck | Cat# S6852 |
| BME | Sigma-Aldrich | Cat# M3418 |
| Recombinant Activin A | R&D system | Cat# 338-AC |
| Recombinant bFGF | R&D system | Cat# 3718-FB |
| Recombinant Mouse LIF Protein | Millipore | Cat# ESG1107 |
| Recombinant Human LIF Protein | Absin Bioscience Inc | Cat# abs04302 |
| N2 Supplement (100X) | Thermo Fisher Scientific | Cat# 17502048 |
| B27 Supplement (50X) | Thermo Fisher Scientific | Cat# 17504044 |
| DMEM/F12 | Thermo Fisher Scientific | Cat# 21103049 |
| Neurobasal | Thermo Fisher Scientific | Cat# 11540566 |
| Knockout™ DMEM | Thermo Fisher Scientific | Cat# 10829018 |
| Knockout™ SR | Thermo Fisher Scientific | Cat# 10828028 |
| mTeSR | STEMCELL Technology | Cat# 85850 |
| Versene | Thermo Fisher Scientific | Cat# 15040066 |
| MEM Non-Essential Amino Acids | Thermo Fisher Scientific | Cat# 11140050 |
| L-Gultamine | Thermo Fisher Scientific | Cat# A2916801 |
| GlutaMAX | Thermo Fisher Scientific | Cat# 35050061 |
| Antibiotic-Antimycotic | Thermo Fisher Scientific | Cat# 15240112 |
| Accutase | Thermo Fisher Scientific | Cat# A1110501 |
| Trypsin-EDTA (0.5%) | Thermo Fisher Scientific | Cat# 15400054 |
| Matrigel Matrix | Corning | Cat# 354277 |
| TrueCut™ Cas9 Protein v2 | Thermo Fisher Scientific | Cat# A36496 |
| Terminal Deoxynucleotidyl Transferase | Thermo Fisher Scientific | Cat# 10533065 |
| dATP | Roche | Cat# 11051440001 |
| **Critical Commercial Assays** | | |
| OxiSelect In Vitro ROS/RNS Assay Kit | Cell Biolabs | Cat# STA-347 |
| XF Cell Mito Stress Test Kit | Seahorse Bioscience | Cat# 103015-100 |
| Pierce BCA Protein Assay Kit | Thermo Fisher Scientific | Cat# 23227 |
| RIPA Lysis and Extraction Buffer | Thermo Fisher Scientific | Cat# 89900 |
| TURBO DNA-free Kit | Thermo Fisher Scientific | Cat# AM1907 |
| TRIzol | Thermo Fisher Scientific | Cat# 15596026 |
| PowerUp™ SYBR™ Green Master Mix | Thermo Fisher Scientific | Cat# A25776 |
| GoTaq® G2 Master Mixes | Promega | Cat# M7823 |
| SuperScript IV First-Strand Synthesis System for RT-PCR | Thermo Fisher Scientific | Cat# 18091050 |
| TransScript® Uni All-in-One First-Strand cDNA Synthesis SuperMix for qPCR (One-Step gDNA Removal) | TranGen Biotech | Cat# AU341 |
| EDTA-free protease inhibitors | Roche | Cat# 11836153001 |
| High-sig ECL Western Blotting Substrate | Tanon | Cat# 180-501 |
| GeneArt Precision gRNA Synthesis Kit | Thermo Fisher Scientific | Cat# A29377 |
| Neon™ Transfection System | Thermo Fisher Scientific | Cat# MPK1025 |
| X-tremeGENE™ HP DNA Transfection Reagent | Roche | Cat# 6366236001 |
| Lipofectamine® 3000 Reagent | Thermo Fisher Scientific | Cat# L3000001 |
| pEASY®-T1 Cloning Kit | TranGen Biotech | Cat# CT101-01 |

| | | |
|---|---|---|
| **Experimental Models: Cell Lines** | | |
| Feeder | This paper | N/A |
| H9 hESC | WiCell | WA09 |
| R1 mESC | ATCC | SCRC-1011 |
| H9-C8 | This paper | N/A |
| H9-5i | This paper | NA |
| H9-MSI1(138-362) | This paper | NA |
| H9-MSI1(272-362) | This paper | NA |
| **Oligonucleotides** | | |
| See Supplementary Table S4 | This paper | N/A |
| **Recombinant DNA** | | |
| CSII-EF-Flag-MSI1-2A-Neo | This paper | N/A |
| CSII-EF-Flag-MSI1(138-362)-2A-Neo | This paper | N/A |
| CSII-EF-Flag-MSI1(272-362)-2A-Neo | This paper | N/A |
| CSII-EF-MSC | This paper | N/A |
| **Software and Algorithms** | | |
| Adobe Photoshop (Adobe) | Adobe Systems | http://www.adobe.com/products/photoshop.html |
| Image J | Fiji | https://imagej.nih.gov/ij/ |
| GraphPad Prism 6 | GraphPad Software, Inc | https://www.graphpad.com/scientific-software/prism/ |
| Excel | Microsoft | https://www.microsoft.com/en-gb/ |
| R v3.6.2 | N/A | https://www.R-project.org/ |
| Trimmomatic | (Bolger *et al*, 2014) | http://www.usadellab.org/cms/uploads/supplementary/Trimmomatic |
| hisat2 | N/A | https://ccb.jhu.edu/software/hisat2/index.shtml |
| qualimap_v2.2.1 | N/A | http://qualimap.bioinfo.cipf.es/ |
| HTSeq | N/A | https://htseq.readthedocs.io/en/release_0.11.1/ |
| edgeR | (Robinson *et al*, 2010) | http://www.bioconductor.org/packages/release/bioc/html/edgeR.html/ |
| Venny | N/A | https://bioinfogp.cnb.csic.es/tools/venny/index.html |
| Panther | (Mi *et al*, 2021) | http://pantherdb.org |
